# Crop-specific agricultural land use and Rift Valley fever outbreaks in Uganda: a longitudinal compositional analysis

**DOI:** 10.64898/2026.06.24.734419

**Authors:** Carson Telford, Luke Nyakarahuka, Jimmy Baluku, Joanita Mutesi, Conghe Song, Ross Boyce, Mike Emch, Jess Edwards, Trevor Shoemaker, Justin Lessler

**Affiliations:** University of North Carolina, Chapel Hill, North Carolina; Uganda Virus Research Institute, Kampala, Uganda; Viral Special Pathogens Branch, Centers for Disease Control and Prevention, Atlanta, Georgia

## Abstract

Rift Valley fever (RVF) is a mosquito-borne disease that can cause severe illness and death in both humans and livestock. Since 2016, Uganda has experienced recurrent but localized RVF outbreaks concentrated in the country’s southwestern region. The ecological drivers of this emergence remain unclear, as outbreaks have occurred throughout the year and show little association with meteorological patterns. We evaluated whether crop cultivation, particularly banana cultivation, is associated with RVF outbreak occurrence after controlling for likely confounders. We conducted a longitudinal study of human-inhabited 5 × 5 km grid cells across southwestern Uganda from 2016–2024. Annual Sentinel-2 satellite imagery composites were used to classify land cover into banana, coffee, ground crops, and non-crop categories, and the proportion of each land type was calculated for every grid-cell year. Because land cover proportions are compositional, isometric log-ratio transformations were used to estimate the independent effects of each land type. Confounding was addressed through propensity weighting, and crop substitution effects were estimated using g-computation. Banana land cover was the only land type consistently associated with increased RVF outbreak likelihood. In grid-cell years with low baseline banana cover, a 10-percentage point substitution from other land classes into banana was associated with a 1.64-fold increase in the odds of an RVF outbreak (95% CI: 1.17–2.29). In a simplified banana-only model, each 10-percentage point increase in banana cover was associated with a 1.21-fold increase in outbreak odds (95% CI: 1.02–1.43). Holding banana cover constant, substitutions among coffee, ground crop, and non-crop land showed weak or null associations. These findings suggest that banana cultivation may be an important ecological feature influencing RVF transmission dynamics and outbreak risk in southwestern Uganda.

**Author Summary:** Rift Valley fever (RVF) is a mosquito-borne disease that affects both humans and livestock and has caused repeated outbreaks in southwestern Uganda since 2016. While rainfall and flooding are often linked to RVF outbreaks elsewhere, Uganda’s recent outbreaks have occurred across seasons and are not well explained by weather patterns alone. We investigated whether agricultural land use could help explain where outbreaks occur. Using satellite imagery from 2016–2024, we measured the amount of banana cultivation, coffee cultivation, ground crops, and non-crop land across southwestern Uganda and evaluated their association with RVF outbreak occurrence. We found that areas with greater banana cultivation were consistently more likely to experience RVF outbreaks, even after accounting for environmental and demographic factors. In contrast, coffee, ground crops, and non-crop land showed little evidence of an independent association with outbreak risk. These findings suggest that banana cultivation may create ecological conditions that favor RVF transmission. Rather than indicating that bananas themselves cause disease, the results point to banana-growing landscapes as potential environments where interactions among mosquitoes, livestock, and humans may increase transmission opportunities. Understanding these local ecological drivers could help improve surveillance, risk assessment, and prevention strategies for RVF in Uganda and other endemic regions.

## Introduction

Rift Valley fever (RVF) is a mosquito-borne zoonotic disease resulting from infection with Rift Valley fever virus (RVFV). RVF can lead to severe, potentially fatal disease in humans and domestic livestock. Due to high mortality among livestock and 100% fetal abortion rates in pregnant livestock, RVF outbreaks can cause considerable economic loss in agriculturally driven economies^1,2^. *Aedes* mosquito species are the primary vectors of RVFV, spreading the virus by biting susceptible humans, domesticated livestock, and wild animals. These *Aedes* species also transmit the virus transovarially to their eggs, which can persist in a desiccated state for years^3,4^. After outbreaks begin, secondary vectors such as *Culex*, *Mansonia*, *Coquillettidia*, and *Eretmapodites* species can amplify transmission^5^. Human infections mostly occur through direct contact with infected animals rather than via mosquito bites, as many of the RVFV vectors are primarily zoophilic^2,5^. Large outbreaks are typically linked to heavy rainfall and flooding, often correlating with the El Niño Southern Oscillation cycle, which can cause mass hatching of desiccated transovarially infected *Aedes* mosquito eggs^3,4,6–8^. During the years between these epidemic events, low level “interepidemic” RVFV circulation occurs but is poorly understood^9–13^.

In Uganda, RVF reemerged in 2016 after over 40 years of absence, with outbreaks occurring almost exclusively in the southwest highland region^13–16^. From 2016-2024, 156 human infections resulting from 90 separate outbreaks (defined as at least 1 case) were identified^13,14^. In contrast to countries such as Kenya or Tanzania which experience periodic RVF epidemics following anomalous precipitation (such as El Nino Southern Oscillation events), these outbreaks had no clear correlation with meteorological patterns and occurred during all months of the year^13,16^.

Most livestock in Uganda are located within the “Cattle Corridor”, a band of land stretching from the southwestern corner of the country to the northeastern corner. A conventional explanation for the increased risk of RVF in the southwestern region is that it has a high density of RVF-susceptible livestock (cattle, sheep, goats), which act as amplifiers for human infection (Telford, Nyakarahuka in press). However, the absence of outbreaks and overall lower seropositivity among livestock in the NE region of the Cattle Corridor suggests that factors in addition to livestock density may drive the spatial distribution of RVFV in Uganda^15^.

Agricultural activities have been shown to be associated with increased prevalence of multiple mosquito species (including *Aedes* species) in other areas of the world ^17–21^. The southwest of Uganda is a major agricultural region, and home to the majority of Uganda’s banana clutivation^22^. *Aedes* mosquitoes have been observed to dominate crop plantation settings relative to other mosquito species, with increased biting surrounding perennial evergreen broadleaf crops, such as banana and coffee, which keep their leaves year-round^23,24^. A study conducted in southwest Uganda observed preferential breeding of *Aedes* mosquitos in the axils of banana plants (where the leaf emerges from the stem)^24^. Banana axils retain moisture and organic matter, providing an ideal environment for *Aedes* mosquito species to lay their eggs, after which subsequent temporary pooling of water within the axils causes eggs to hatch. Further, crop cultivation and livestock ownership often co-occur. Hence, we hypothesize that banana cultivation is a critical driver of the RVFV transmission cycle in Uganda, and responsible for the year-round circulation and spatial distribution of RVF in the country.

To explore this hypothesis, we here present a longitudinal study of RVF outbreaks in southwest Uganda (2016–2024), where we analyze the association between cultivation of specific common crop types in Uganda and the likelihood of a human RVF outbreak, adjusting for key confounders of this relationship.

## Methods

### Ethical Statement

RVF outbreak data were derived from investigations conducted by the Uganda Ministry of Health, Uganda Virus Research Institute (UVRI), and collaborating partners as part of routine outbreak response activities. These activities were conducted under public health response authorities and were determined by the U.S. Centers for Disease Control and Prevention to be non-research public health practice. Use of these de-identified data for analysis presented here was approved by the University of North Carolina Institutional Review Board and the UVRI Research Ethics Committee (UVRI REC: GC/127/16/03/551).

### Study design

The units of analysis for this study were 5x5-kilometer grid cells with evidence of human habitation (as assessed by Center for International Earth Science Information Network of Columbia University ^25^), covering the southwestern region of Uganda (approximately 104,000 km^2^). A grid-cell resolution of 5x5-kilometers was selected as it approximates maximum livestock grazing ranges^15^. Each grid cell was analyzed annually from 2016-2024. We excluded grid cells with >90% water cover to ensure stability in land cover proportion calculations. Overall, there were 3,012 grid cells included, resulting in a total of 27,108 grid-cell years.

### Outcome measures

The outcome was whether one or more confirmed RVF cases occurred in a grid cell during a given year. Identification of RVF cases occurs through Uganda National Viral Hemorrhagic Fever surveillance system managed by UVRI and the Uganda Ministry of Health^26^. If physicians at surveillance sites suspected infection with a viral hemorrhagic fever, a sample was collected and sent for confirmatory testing at UVRI^27^. For RVF-positive samples, in-depth investigations were conducted by UVRI, during which geocoordinates were recorded for the residence of confirmed cases^13,14,26^. Over the study period, 90% (90/100) of confirmed human outbreaks in Uganda occurred within the southwest region. Each outbreak was assigned to the grid-cell containing the case household during the year in which it occurred.

### Quantifying exposures through land type classification

The exposures of interest in this study were the proportions of land area classified as banana, coffee, ground crops, or non-crop land per grid-cell year, which summed to 100% of land area in each grid cell.

To quantify these proportions, a land type classification analysis was done using publicly available satellite imagery from the Copernicus Sentinel-2 dataset on Google Earth Engine (GEE)^28,29^. Sentinel-2 captures multispectral reflectance across 13 spectral bands at a 10-meter resolution, including visible, red-edge, near-infrared, and shortwave-infrared wavelengths, with revisit times of approximately five days^29^. We created annual median composites of imagery corresponding to each year from 2016-2024 within our study area. Cloud-contaminated observations were removed prior to compositing, and annual median composites were generated from the remaining cloud-free observations available throughout each year. Composite annual images were resampled with bilinear mean interpolation to a spatial resolution of 20-meters. Additional bands were computed for the normalized difference vegetation index (NDVI) (a measure of vegetation greenness) and the Red Edge 1 band (a Sentinel-2 band that is particularly sensitive to vegetation shifts)^30,31^. These included annual average NDVI, NDVI during the wet season months, NDVI during the dry season months, NDVI standard deviation, Red Edge 1 during the wet season, Red Edge 1 during the dry season, and Red Edge standard deviation. Bands aggregated to spatial resolutions of 60- and 100-meters were also added for the NDVI and Red Edge 1 variables to capture neighborhood context. Full details on the 24 features extracted are available in the supplemental material.

We developed a human-verified training dataset to support development of a pixel-level land type classifier that assigned pixels to one of eight categories: banana, coffee, ground crops, grass, bush, forest, urban, and water. To facilitate accurate visual identification, UVRI provided five reference coordinates for each land class. These points were used as references to train a human classifier to distinguish land use categories in yearly ultra-high resolution (30 centimeters) Google Earth images (range 2016-2024). A random sample of one percent of 1 x 1 kilometer grid cells from the study area was selected for manual annotation. A single analyst drew polygons corresponding to land use classes within Google Earth, thereby generating year-specific labeled datasets. The annotation process began with 2024, which served as the baseline dataset. This dataset was then overlaid onto imagery from the preceding year. Where discrepancies in land use classification were identified, new polygons were delineated and assigned the appropriate labels. This iterative procedure was repeated sequentially for each year back to 2016. If land use within a polygon area could not be determined due to unavailable or poor-quality Google Earth imagery during a given year, the polygon was omitted for that year.

The final training samples consisted of a set of polygons for each year in the study period. Training assignments were compared with two standard crop classification datasets (MODIS Land Cover and Dynamic World) and a subset of discrepancies were explored. The baseline training dataset that labeled land cover in 2024 included 4,371 polygons, representing a cumulative area of 735.33 km^2^ (the total study area represented an area of 75,960 km^2^). The relative proportion of polygons corresponding to each land cover class and the average area covered by each polygon class was consistent across years (Table S1). Labeled pixel values were extracted into a tabular dataset with a row for each pixel and columns for the satellite imagery features (as described above) and the land type label. This dataset was randomly partitioned into training (70%) and test (30%) subsets with clustering at the 5x5 km study grid cell level to prevent information leakage between the train and test sets.

Individual classification models were trained separately for each year using gradient-boosted decision trees implemented in the XGBoost package within R^32,33^. This year-specific modeling approach was used to maximize prediction performance by mitigating inconsistent spectral signals arising from inter-annual variation in climate and satellite image availability. For each annual dataset, we tuned the following hyperparameters: number of trees, maximum tree depth, learning rate, subsample fraction of observations, and the proportion of features randomly sampled for each split. Hyperparameter selection was conducted via a predefined grid search combined with spatially clustered five-fold cross-validation to account for spatial autocorrelation in the training data. Models were trained using a multi-class softmax cross-entropy loss function. To address class imbalance, inverse-frequency class weights were applied during training.

Hyperparameter selection was based on maximizing the product of sensitivity and specificity for the pooled crop classes (banana, coffee, and ground crops) versus non-crop classes via cross-validation. Cross-validation was performed using five spatially clustered folds, created by partitioning the 5 × 5 km study grid cells containing labeled training pixels into five geographically distinct groups. During each iteration, models were trained on four folds and evaluated on the remaining fold to account for spatial autocorrelation and provide a more realistic assessment of predictive performance.

The final model was evaluated using cross-validation performance within the training dataset and predictive performance of the fully trained model on the held-out test dataset. Because the primary objective was accurate quantification of crop exposures, evaluation focused on class-specific measures of crop classification performance. Performance summary metrics included recall (sensitivity, equivalent to producer’s accuracy), precision (positive predicted value [PPV], equivalent to user’s accuracy), and specificity. Overall classification accuracy was also reported across the four exposure classes (banana, coffee, ground crops, and non-crop).Classifier training was assessed using calibration plots for each crop type, examining the relationship between predicted probabilities of class membership and the observed proportion of pixels belonging to that class.

Final tuned models for each year were fit using all available labeled data and applied to classify the land type of all pixels in the full study area from 2016-2024 (original human-classified polygons were ignored). The resulting classified rasters were used to compute the proportion of land area classified as banana, coffee, ground crops, and non-crop (which encompassed grass, bush, forest, or urban) in each 5 × 5 km grid-cell year. Pixels classified as water were excluded from these calculations.

### Identification and collection of key covariates

To identify what additional data needed to be collected, we first encoded causal relationships between key variables that might be associated with study observations or outcomes in a directed acyclic graph (DAG). The DAG was then analyzed to identify potential confounders and appropriate adjustments to reduce confounding (Figure S1)^34^. Based on this analysis, we identified key variables for collection and adjustment: precipitation patterns, human and livestock population density, elevation-derived topographic measures, and soil texture characteristics. Covariates were obtained from publicly available gridded raster datasets (see Supplementary Methods for details) and summarized within each 5 × 5 km study grid cell-year. A descriptive analysis compared covariate values across outbreak and non-outbreak grid cells using a 2-sample t-test with an alpha value of 0.05.

### Compositional data analysis

Proportions of banana, coffee, ground crops, and non-crop land are dependent, such that an increase in one class implies a decrease in at least one other class. To address this, we applied isometric log-ratio (ILR) transformations to the four land classes with sequential binary partitions, which resulted in 3 distinct ILR variables^35–38^. ILR transformation converts the proportions into variables that reflect relative differences between land types (i.e., how one class compares to others), rather than their raw percentages. This removes the constraint that values must sum to 100% and allows the variables to be analyzed independently. By modeling ILR-transformed variables, we account for the relative nature of land cover while enabling valid use of standard statistical methods.

The marginal effects of the ILR terms on RVF outbreak likelihood were estimated in a single model using binomial generalized estimating equations (GEE). This model incorporated propensity weights to control for confounding (see weighting methods below) and clustering at the 5x5 km grid-cell level to account for repeated annual observations. Fitted ILR coefficients can be thought of as representing how RVF outbreak likelihood changes as land cover shifts from one set of classes toward another within a given ILR balance. Positive and negative coefficients indicate whether a relative shift was positively or negatively associated with RVF outbreak likelihood. We used g-computation to transform fitted ILR coefficients into more interpretable effect estimates^39^. Specifically, for each grid-cell year measured in our dataset, we applied reallocations (substitutions) of land cover proportions (1–30%) between banana, coffee, ground crops, and non-crop classes. ILR transformations were then applied to these land cover compositions. The fitted model was then used to predict the log-odds of an RVF outbreak resulting from the observed and counterfactual compositions. Odds ratios were derived as the contrast between the observed and counterfactual log odds estimates. Confidence intervals for g-computation-derived odds ratios were calculated using the robust covariance matrix from the fitted GEE model. Standard errors were estimated from the change in ILR-transformed land-cover composition between the observed and counterfactual scenarios. Confidence intervals were constructed on the log-odds-ratio scale and exponentiated to obtain odds-ratio intervals.

We evaluated how RVF outbreak likelihood changed under two types of land-cover reallocations. First, we evaluated pairwise substitutions between two classes, holding the remaining two constant (e.g., increasing banana cover by 10% while decreasing coffee cover by 10%). Second, we evaluated proportional reallocations in which increases in one class were drawn proportionally from the remaining three classes combined (e.g., increasing banana cover by 10% through proportional reductions in coffee, ground crops, and non-crop land). Effects were summarized across substitution magnitudes (1–30%) and stratified by baseline abundance of the receiving class (low: <25th percentile, moderate: 25th–75th percentile, high: >75th percentile). Counterfactual scenarios were only evaluated when sufficient land cover was available in the source class(es) to support the specified reallocation magnitude. For example, a grid-cell containing 10% banana cover could not be evaluated under a 30% reallocation from banana to another class.

### Propensity weighting

Propensity weights to control for confounding were derived from the modeled joint inverse density across the 3 ILR terms. A weighting approach was selected because only 90 RVF outbreaks were observed during the study period, whereas adjustment was required for numerous potential confounders. Estimating confounding through the propensity model allowed adjustment to be based on the full cohort while preserving a parsimonious outcome model focused on estimating the marginal association between land-cover composition and RVF outbreak likelihood.

To do this, we first fit separate propensity models for each ILR term conditional on other covariates^40^. To capture nonlinearities a Super Learner ensemble to predict the continuous ILR outcome was fit that included linear regression, generalized additive models, and random forests^41^. The joint conditional density produced by each of the three ILR-specific Super Learner ensembles was modeled as a multivariate normal distribution with mean given by Super Learner predictions for each ILR term and covariance estimated from the residuals of each model (assuming independent variances)^40^. Stabilized inverse density (propensity) weights were constructed as the ratio of the marginal joint density of the ILR components to their conditional joint density given covariates. Weights were truncated at the 1st and 99th percentiles to reduce sensitivity to extreme values.

Covariate balance across levels of the ILR terms was assessed pre- and post- application of propensity weights using correlation coefficients and standardized mean differences (SMDs)^42^. SMDs were defined as standardized differences between covariate means within quintiles of each ILR term and the overall sample mean. A threshold of 0.2 was used to define acceptable covariate balance^43^.

### Simplified banana-only model

Results from the compositional data analysis highlighted banana crops as the single land type that was consistently associated with RVF outbreaks. To provide a more interpretable effect estimate, we fit a logistic regression model of effect of a 10-unit increase in the proportion of banana-cultivated land area on RVF incidence. Confounding was addressed using the same propensity weighting approach as described above, with weights estimated directly for the proportion of banana land cover.

### Sensitivity analysis

We evaluated the robustness of our findings to uncertainty in land type classifications. All pixels’ landcover classes were resampled based upon predicted class probabilities (from the landcover classification algorithm) 1000 times, and the full estimation procedure was run on each resampled dataset. This resulted in 1000 individual effect estimates for both the compositional and simplified banana-only analyses. The distribution of estimates (and implied substitution effect) was compared to estimates from the main analysis.

## Results

### Land cover classification

The land cover classifier demonstrated strong predictive performance across major land-cover classes when evaluated on the held-out test data set. For bananas, average sensitivity was 0.92, specificity was 0.99, and positive predictive value (PPV) was 0.91. For coffee, average sensitivity was 0.86, specificity was 0.99, and PPV was 0.92. For ground crops, average sensitivity was 0.77, specificity was 0.99, and PPV was 0.83 (Figures S2–S3). Overall classification accuracy across banana, coffee, ground crops, non-crop land classes, and water was 0.93 (Table S2). Classification performance was similar in cross-validation and held-out test datasets.

Most classification error involved confusion between banana and ground crops. Calibration analyses demonstrated good agreement between predicted probabilities and observed class frequencies, and the Uganda-specific classifier outperformed publicly available land-cover products when evaluated against independently labeled test data. Additional validation results are provided in the Supplementary Materials (Figures S2–S9). The spatial distribution of modal land-cover classes from 2016–2024 is shown in Figure 1.

**Figure 1.**
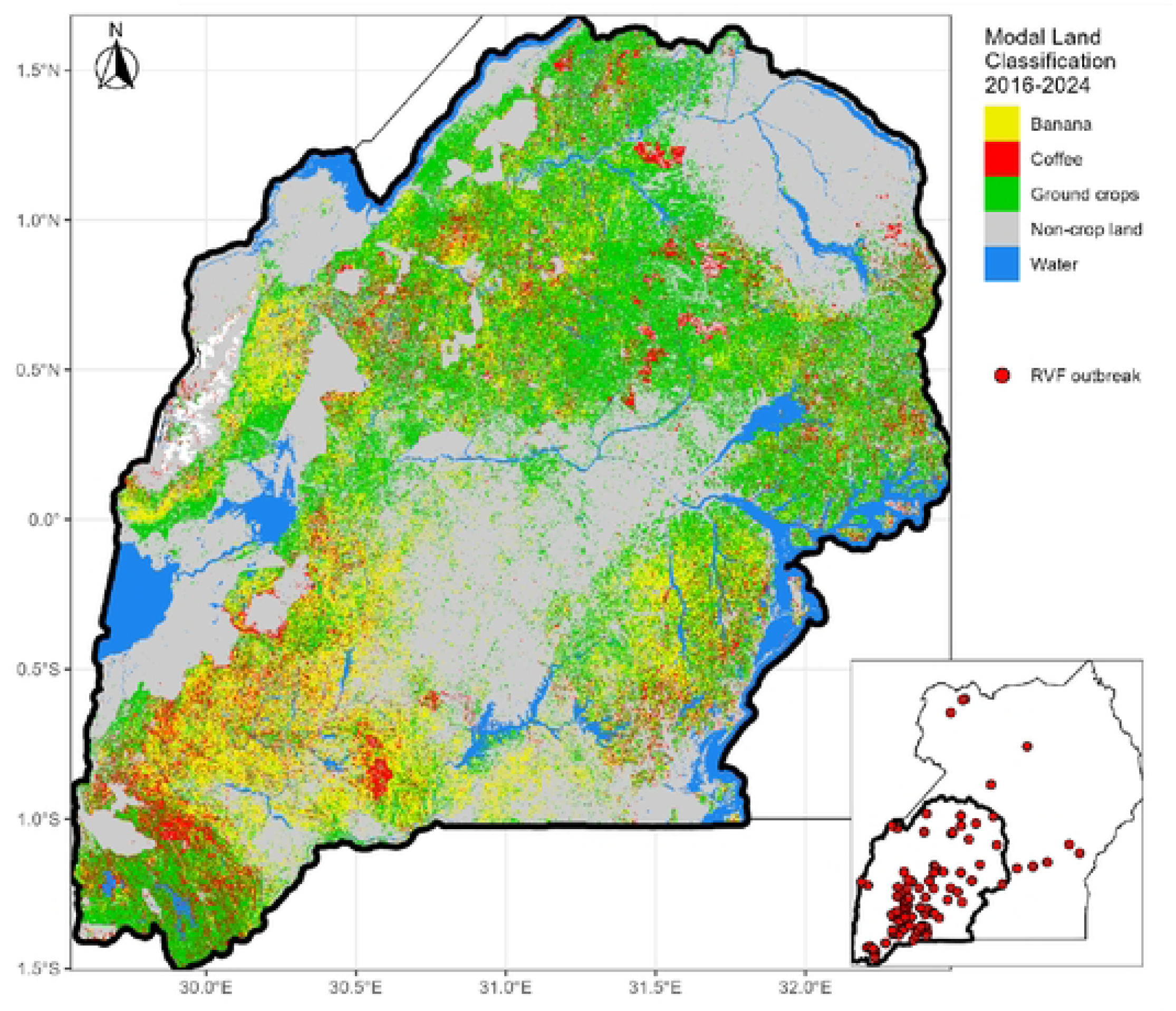
Land type classification (modal class) and locations ofRVF outbreaks in southwest Uganda fro1n 2016-2024, Non-crop land is coinprised of grass, bush, forest, and urban land classes.

### Descriptive analysis

Across 5×5 km grid cells from 2016–2024, non-crop land accounted for the largest proportion of land area on average (0.51), followed by ground crops (0.27), banana (0.14), and coffee (0.09). The crude mean proportion of banana land cover was higher in grid cells with an RVF outbreak (0.23) compared to those without an outbreak (0.14) (p<0.001). In contrast, ground crops (p=0.0576) and non-crop land (p=0.0678) proportions tended to be lower in outbreak grid cells compared to non-outbreak grid cells. The proportion of coffee land area was similar between outbreak (10%) and non-outbreak grid cell years (9%) (p=0.27) (Table 1). Grid cell-years with RVF outbreaks were also characterized by statistically higher mean human and livestock densities, greater within-year precipitation variability (coefficient of variation), and higher elevation. Total annual precipitation was lower on average at outbreak grid cells compared to non-outbreak grid cells. Unadjusted analysis did not find evidence of statistically significant differences in TWI, precipitation anomalies, slope, or the proportion of clay soil content (Table 1).

**Table 1.** Mean distributions of land cover compositional proportions and confounding variables among 5 x 5-kilometer grid-cells in Southwestern Uganda measured annually from 2016-2024.

|  | No Outbreak<br>(N=27018)<br>*Mean (SD)* | RVF Outbreak<br>(N=90)<br>*Mean (SD)* | T-test<br>*p-value* |
| --- | --- | --- | --- |
| **Land Cover Proportion** |  |  |  |
| Pooled crops | 0.49 (± 0.32) | 0.55 (± 0.30) | 0.0678 |
| Non-crop land | 0.51 (± 0.32) | 0.45 (± 0.30) | 0.0678 |
| Banana | 0.14 (± 0.15) | 0.23 (± 0.19) | <0.001 |
| Coffee | 0.086 (± 0.097) | 0.099 (± 0.11) | 0.272 |
| Ground crops | 0.27 (± 0.22) | 0.23 (± 0.21) | 0.0576 |
| **Population measures** |  |  |  |
| Human population density* | 190 (± 170) | 280 (± 270) | 0.00438 |
| Livestock population density** | 4100 (± 3000) | 5400 (± 2800) | <0.001 |
| **Climate measures** |  |  |  |
| Annual precipitation (mm) | 1200 (± 190) | 1100 (± 150) | <0.001 |
| Annual coefficient of variation of monthly precipitation (mm) | 1.7 (± 0.22) | 1.9 (± 0.28) | <0.001 |
| Standardized annual precipitation anomaly (z-score) | 0 (± 0.94) | 0.084 (± 0.83) | 0.336 |
| **Topography** |  |  |  |
| Elevation (m) | 1300 (± 290) | 1400 (± 260) | <0.001 |
| Slope (degrees) | 5.3 (± 4.2) | 6.0 (± 3.8) | 0.082 |
| Topographic wetness index | 6.8 (± 3.4) | 6.6 (± 2.8) | 0.371 |
| Clay % | 38 (± 3.2) | 38 (± 3.1) | 0.596 |
| *Persons per square kilometer |  |  |  |
| **Livestock per square kilometer |  |  |  |

### Propensity weighting

Propensity weighting substantially improved covariate balance between compositional land-cover variables and measured confounders. Stabilized joint density weights ranged from 0.05–9.67, corresponding to an effective sample size of 5,827. Correlations between isometric log-ratio (ILR) land-cover variables and measured confounders were reduced after weighting, particularly for ILR1 (banana versus all other land classes) and ILR2 (coffee versus remaining land classes) (Figure S10).

Covariate balance assessed using standardized mean differences (SMDs) also improved after weighting. Balance improvement was greatest for ILR1 and ILR2, with most weighted comparisons meeting or approaching the prespecified SMD threshold of 0.2. Residual imbalance remained for ILR3 (ground crops versus non-crop land) despite weighting (Figure S11). Overall, weighting substantially improved balance for the primary banana and coffee land-cover contrasts evaluated in subsequent analyses.

### Land type reallocation effects

Substitutions of land cover toward banana cultivation were consistently associated with higher odds of RVF outbreaks. The magnitude of this effect was strongest in areas with low baseline banana cover and attenuated as baseline banana cover increased. In low banana-cover areas, a 10% substitution from non-crop land to banana was associated with an average OR of 1.66 (95% CI: 1.27–2.16). The average OR for substitution from coffee to banana was 1.65 (95% CI: 1.05–2.61). The average OR for substitution from ground crops to banana was 1.54 (95% CI: 1.14–2.08). A proportional 10% substitution from all other land classes (coffee, ground crops, non-crop land) to banana produced a similar estimate (OR: 1.64; 95% CI: 1.17–2.29) (Figure 2).

**Figure 2.**
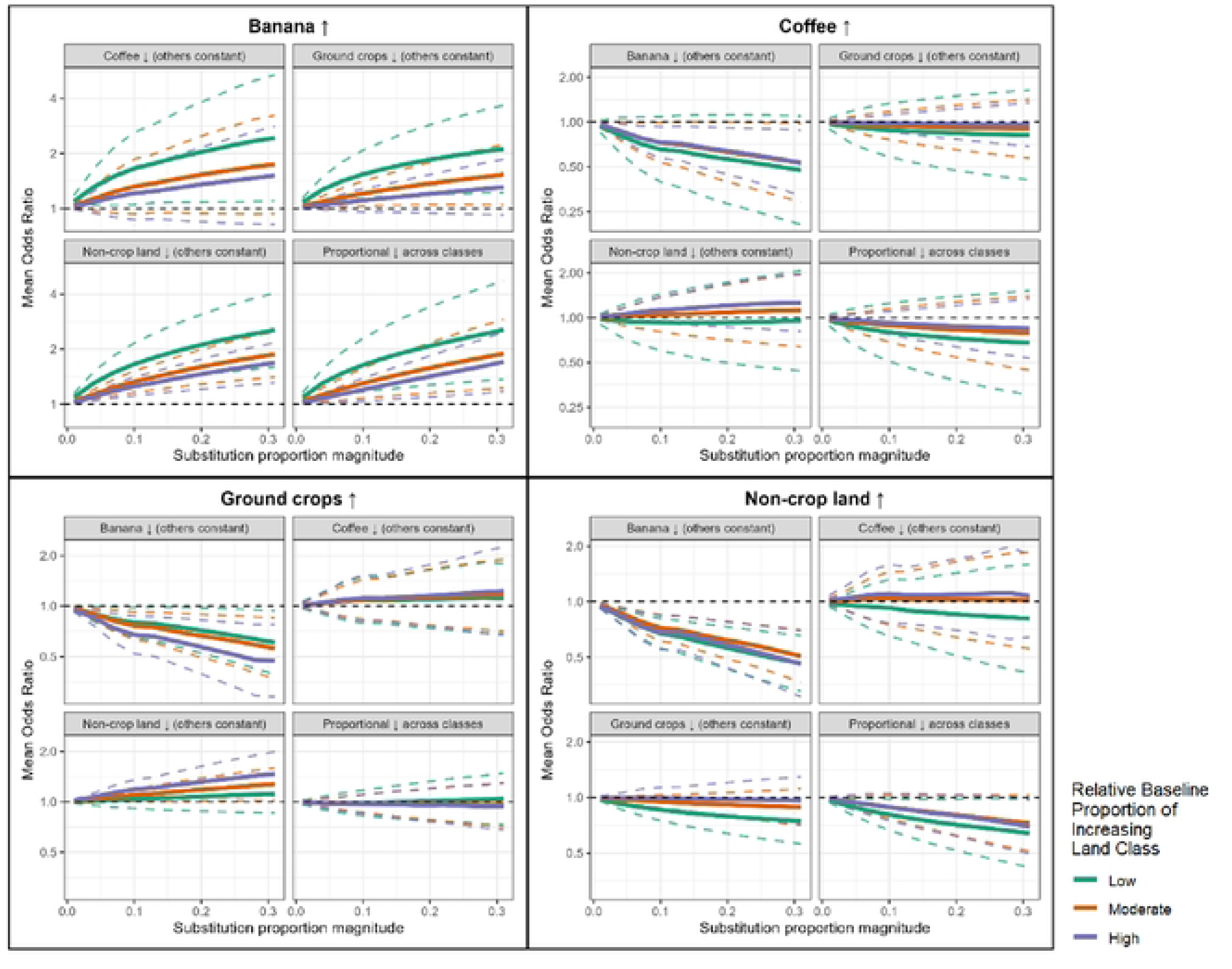
Odds ratio estimates for the substitution effects between banana, coffee, ground crops, and non-crop land across varying levels of baseline land cover proportion and substitution magnitudes.

The magnitude of associations was attenuated in areas with high baseline banana cover. In these areas, a 10% substitution from non-crop land to banana was associated with an average OR of 1.25 (95% CI: 1.11–1.41), while substitution from coffee and ground crops to banana yielded ORs of 1.21 (95% CI: 0.87–1.67) and 1.11 (95% CI: 0.95–1.28), respectively. Larger land-cover substitutions corresponded to larger estimated effects. For example, in areas with moderate banana cover, 10%, 20%, and 30% substitutions from non-crop land to banana were associated with average ORs of 1.33 (95% CI: 1.16–1.52), 1.57 (95% CI: 1.28–1.94), and 1.88 (95% CI: 1.41–2.50), respectively.

Substitutions away from banana cultivation showed the opposite pattern and were generally associated with lower RVF outbreak odds. Increasing coffee at the expense of banana reduced outbreak likelihood, particularly in areas with high baseline coffee cover, where a 10% substitution from banana to coffee corresponded to an average OR of 0.73 (95% CI: 0.57–0.93). Larger substitutions were associated with further decreases in outbreak odds (20%: OR = 0.65, 95% CI: 0.46–0.92; 30%: OR = 0.53, 95% CI: 0.32–0.88). Similar patterns were observed for ground crops. In areas with high baseline ground crop cover, 10%, 20%, and 30% substitutions from banana to ground crops were associated with average ORs of 0.67 (95% CI: 0.52–0.87), 0.58 (95% CI: 0.41–0.83), and 0.47 (95% CI: 0.29–0.79), respectively.

In contrast, substitutions among coffee, ground crops, and non-crop land showed limited evidence of meaningful associations with RVF outbreak likelihood when banana cover was held constant. The main exception was a modest increase in outbreak odds when replacing non-crop land with ground crops (OR: 1.10, 95% CI: 1.01–1.20) (Figure 2).

### Banana-only model estimate

To provide a simplified and readily interpretable estimate of the association between banana cultivation and RVF outbreak occurrence, we fit a model comparing banana land cover to all other land-cover classes combined. Consistent with the compositional analysis, a 10–percentage-point increase in banana cover was associated with 1.21 times higher odds of an RVF outbreak (OR: 1.21, 95% CI: 1.02–1.43). This association was robust to probabilistic reclassification, with sensitivity-analysis estimates ranging from 1.253–1.262 and confidence intervals consistently excluding the null (Figure 3).

**Figure 3.**
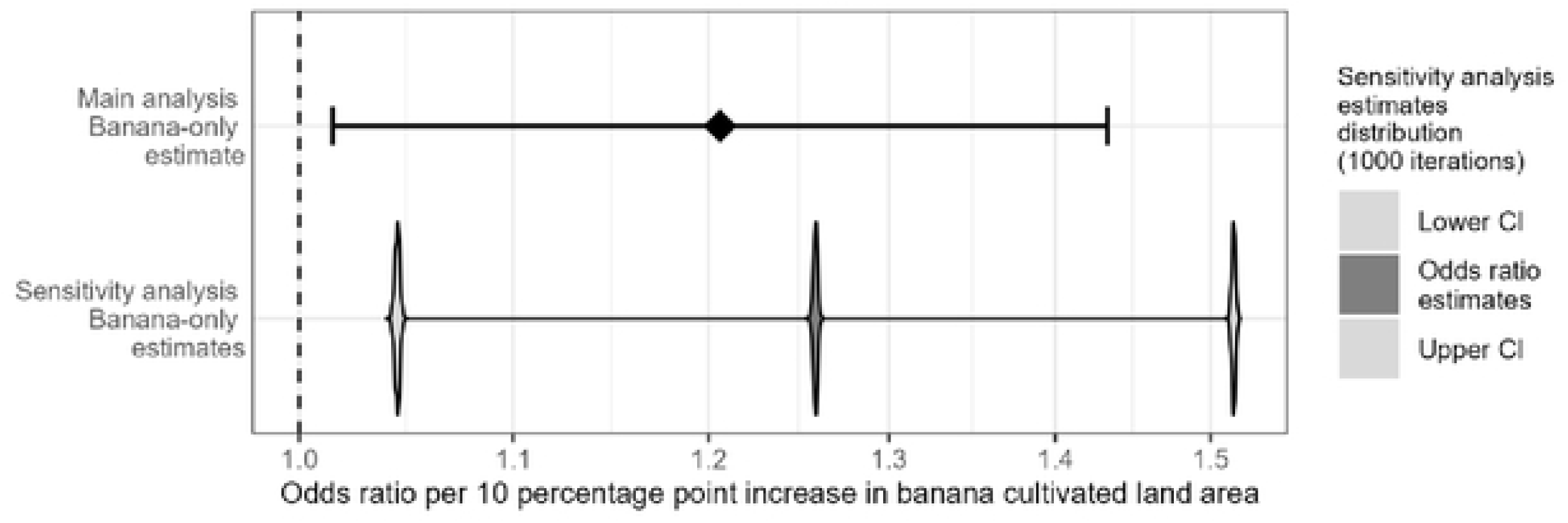
Odds ratio estimate for the average effect of a I0% increase in banana land area relative to all other land area resulting from the main analysis (top), and violin plots visualizing the density of odds ratio estimates with corresponding lower and upper 95% confidence interval bounds resulting from the sensitivity analysis (bottom).

### Probabilistic reclassification sensitivity analysis

Probabilistic reclassification had minimal impact on aggregated land-cover proportions at the 5 × 5 km grid-cell level, despite some variability in classification of individual pixels (Table S3; Figure S12).

Effect estimates from the compositional model were generally robust to classification uncertainty. Estimates involving banana, coffee, and non-crop land were largely unchanged across 1,000 probabilistic reclassification iterations, while effects involving ground crops were attenuated toward the null (Figure S13).

## Discussion

In this longitudinal analysis of RVF outbreaks in southwest Uganda, we found that land-cover composition was associated with RVF outbreak likelihood after adjustment for confounding, with the clearest and most consistent associations observed for banana cultivation. The strongest estimated effects occurred in areas with low baseline banana cover, with a 10% transfer to banana cultivation from other land uses resulted in an increase in RVF risk of over 1.5 times. In contrast, when banana cover was held constant, we found limited evidence that reallocations among coffee, ground crops, and non-crop land were meaningfully associated with RVF outbreak likelihood. Based on these results, we concluded that banana cultivation alone was the primary land use driver of RVF risk, and developed a model of the association between banana crop cultivation and this risk.

Together with prior evidence that *Aedes* mosquitoes breed preferentially in banana plant axils, our findings suggest that banana cultivation may help explain several distinctive features of RVF epidemiology in Uganda that are not well understood^23,24,44^. First, banana axils may support year-round RVFV transmission with limited seasonality, because even small amounts of precipitation may be sufficient to fill axils and trigger hatching of desiccated *Aedes* mosquito eggs. Second, this mechanism may help explain why Ugandan RVF outbreaks are geographically concentrated in the southwest highland region, where banana cultivation is most prevalent, despite high densities of susceptible livestock outside this region^13^. Third, breeding in banana axils may promote small-scale, localized mosquito emergence through both population growth and the hatching of transovarially infected eggs. This localized emergence could help explain the relatively limited size of RVF outbreaks observed in Uganda compared with settings where outbreaks are driven by large-scale flooding events^4,7^.

These findings suggest that RVF risk may be shaped not only by broad climatological anomalies, but also by crop-specific microhabitats that support vector emergence. Although our study focused on southwest Uganda, the co-occurrence of banana cultivation and *Aedes* mosquito species is not unique to Uganda, and similar crop-specific ecological drivers of RVF risk may occur in other banana-growing regions where RVFV is present or could emerge^44^. Even in settings where major outbreaks are typically linked to large-scale flooding or anomalous rainfall, small-scale water pooling associated with specific crops or land management practices may contribute to RVFV persistence and low-level transmission during interepidemic periods. In the case of banana cultivation, risk likely arises from localized, persistent water pools in plant axils that support small-scale Aedes mosquito breeding, rather than from the banana crop itself. Other crops or land management practices that generate standing water and provide suitable microhabitats for mosquito eggs and larvae, whether through rainfall or irrigation, could create comparable ecological niches for vector emergence. By contrast, coffee is often grown on sloped terrain where water is less likely to accumulate, which is consistent with our finding that coffee cultivation was associated with lower RVF risk relative to banana^45^. Together, these considerations suggest that crop-specific water retention and microhabitat formation may be important determinants of RVF risk and interepidemic virus maintenance, alongside established factors such as livestock and wildlife density, climate, and soil characteristics^10^.

These findings suggest that crop-specific land-use information could help spatially target RVF prevention and control strategies in Uganda and similar settings. Identifying banana cultivation as a land-use pattern associated with increased RVF outbreak risk suggests that vector control efforts could be prioritized in areas surrounding banana plantations, particularly in regions with recurrent outbreaks. Targeted larval control or insecticide application around banana plants may be a feasible intervention for government programs or individual farmers, although implementation would require evaluation of effectiveness, cost, and acceptability in local agricultural settings^46,47^. These findings may also inform risk communication and education efforts for individuals with frequent livestock contact, such as herdsmen, who play a central role in managing animal movement and grazing patterns^48,49^. For example, limiting livestock grazing near banana plantations, especially after rainfall, may reduce opportunities for mosquito–livestock contact and subsequent RVFV amplification. More broadly, incorporating crop-specific land-use information into RVF surveillance and early warning systems could improve the spatial targeting of prevention efforts in Uganda without relying solely on acute climatic signals, which appear to be less predictive of outbreak risk in this setting^13^.

The probabilistic classification sensitivity analysis reinforced the interpretation that banana cultivation was the primary crop-specific land-cover feature associated with RVF outbreak risk. Estimated associations for banana were stronger in the sensitivity analysis than in the main analysis, whereas associations for ground crops were attenuated toward the null. In the main analysis, ground crops were marginally associated with outbreak likelihood when replacing non-crop land or coffee, but these associations weakened under probabilistic reclassification. Because probabilistic sampling allowed lower-confidence pixels to switch classes across iterations rather than assigning each pixel deterministically to its most probable class, this attenuation likely reflects uncertainty in ground crop classification. Discordance among crop classes was highest between banana and ground crops, suggesting that part of the ground crop association in the main analysis may have reflected misclassification between these classes (Figure S8). Ground crops also showed the highest discordance with non-crop land, further supporting the interpretation that ground crop estimates were more sensitive to classification uncertainty.

Several limitations should be considered when interpreting these findings. First, our analysis was conducted using 5 × 5 km grid cells, which capture regional land-use patterns rather than fine-scale farm- or herd-level processes. Livestock density, grazing behavior, and proximity to cultivated land may vary substantially within grid cells and could modify the relationship between crop cultivation and RVF risk. Because fine-scale cohort data on livestock herds and farms were not available across the full study area, we used livestock density estimates at 10 × 10 km resolution to control for potential confounding by livestock abundance. Future farm-level studies could help clarify how local crop cultivation patterns, livestock density, and grazing practices jointly shape RVF outbreak risk. Second, several confounders in this analysis, including precipitation, soil, and topography, were derived from satellite-based data products.

Because land cover was also classified using satellite imagery, interpretation of these results assumes that measurement error in land cover classification is sufficiently independent of measurement error in the adjustment variables. Shared environmental information across satellite-derived products could introduce bias if adjustment variables capture features that are also related to land-cover classification. For example, adjustment for environmental variables correlated with crop cultivation could partially attenuate associations if those variables lie along pathways linking crop-specific microhabitats to RVF risk. The direction and magnitude of this potential bias are uncertain and may differ across land-cover classes. Third, inverse density weighting improved covariate balance for most land-cover contrasts, but balance was less complete for ILR3, which represented the log ratio between ground crops and non-crop land.

Ground crops were more prevalent than banana or coffee across the study area and were more strongly correlated with human population density, which may have limited the ability of the weights to balance confounders across ground crop levels without inducing weight instability. As a result, estimates specific to ground crops may be more vulnerable to residual confounding and should be interpreted with caution. Finally, this was an observational study, and unmeasured confounding cannot be fully excluded despite adjustment for potential confounders identified in the causal DAG. However, the consistency of the banana association across compositional reallocation scenarios, the robustness of this association in the probabilistic classification sensitivity analysis, and the biological plausibility of banana plant axils as Aedes mosquito breeding sites support our main conclusion that banana cultivation is an important land-cover feature associated with RVF outbreak likelihood in southwest Uganda.

Overall, these findings support our hypothesis that banana cultivation may drive RVF outbreak likelihood in southwest Uganda. More broadly, they suggest that agricultural land-use patterns—and the crop-specific microhabitats they create—may contribute to RVFV transmission and represent an important target for geographically focused surveillance and prevention strategies.

---

**Figure S1.**
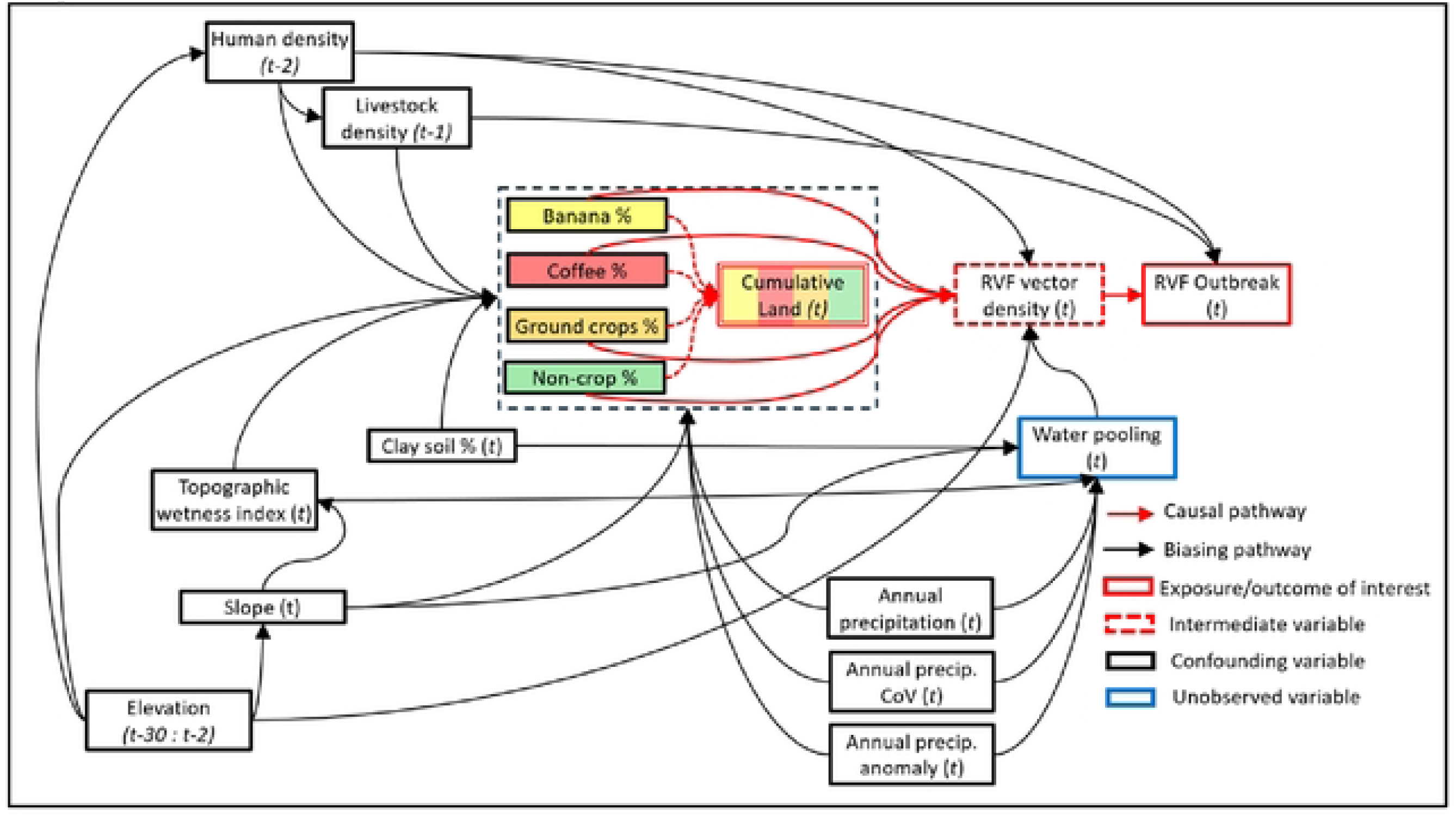
Causal directed acyclic graph depicting the hypothesized causal relationship between crop cultivation and RVF outbreaks.

**Figure S2.**
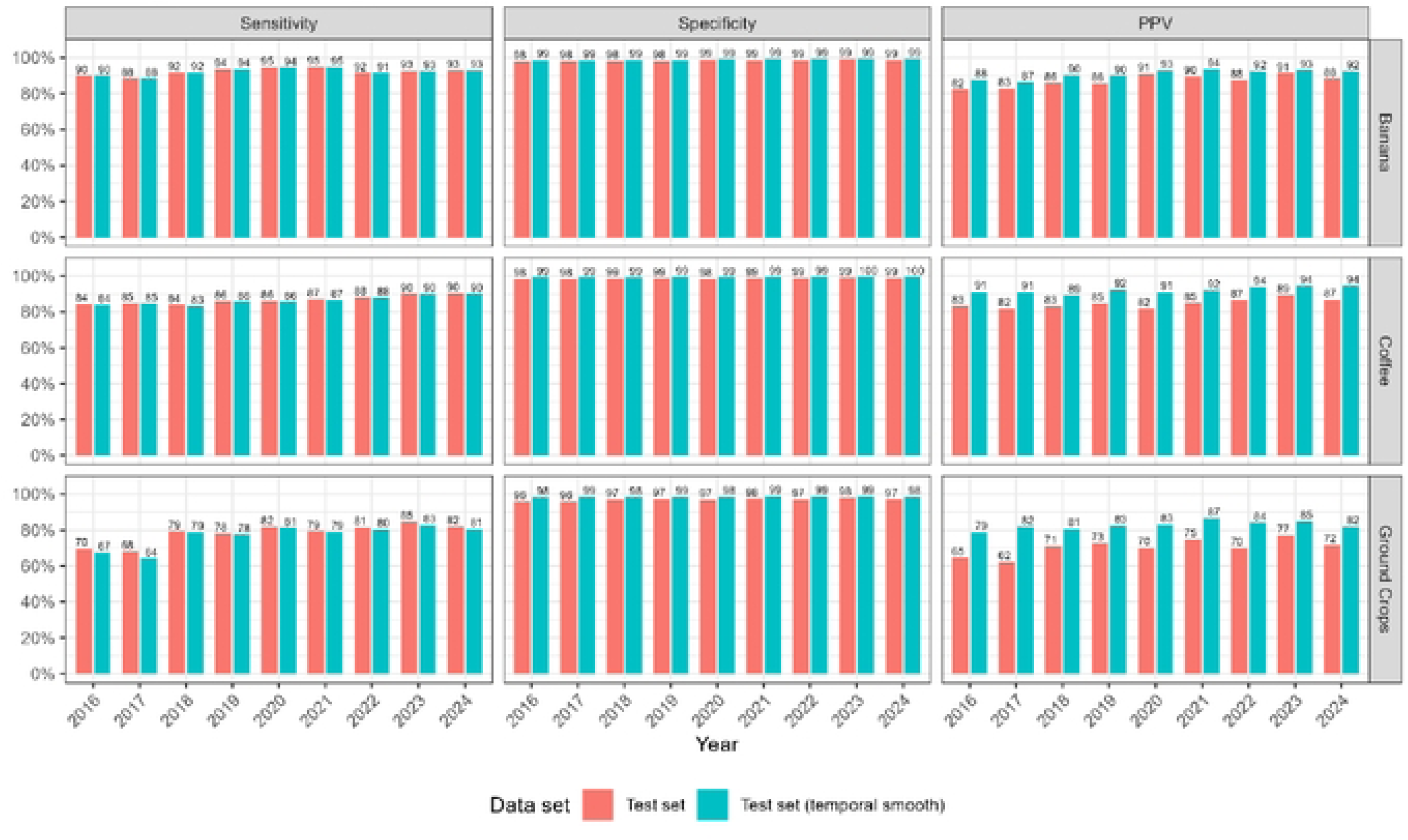
Annual classification performance for crop classes resulting from cross validation on the training data set and prediction on the holdout test set.

**Figure S3.**
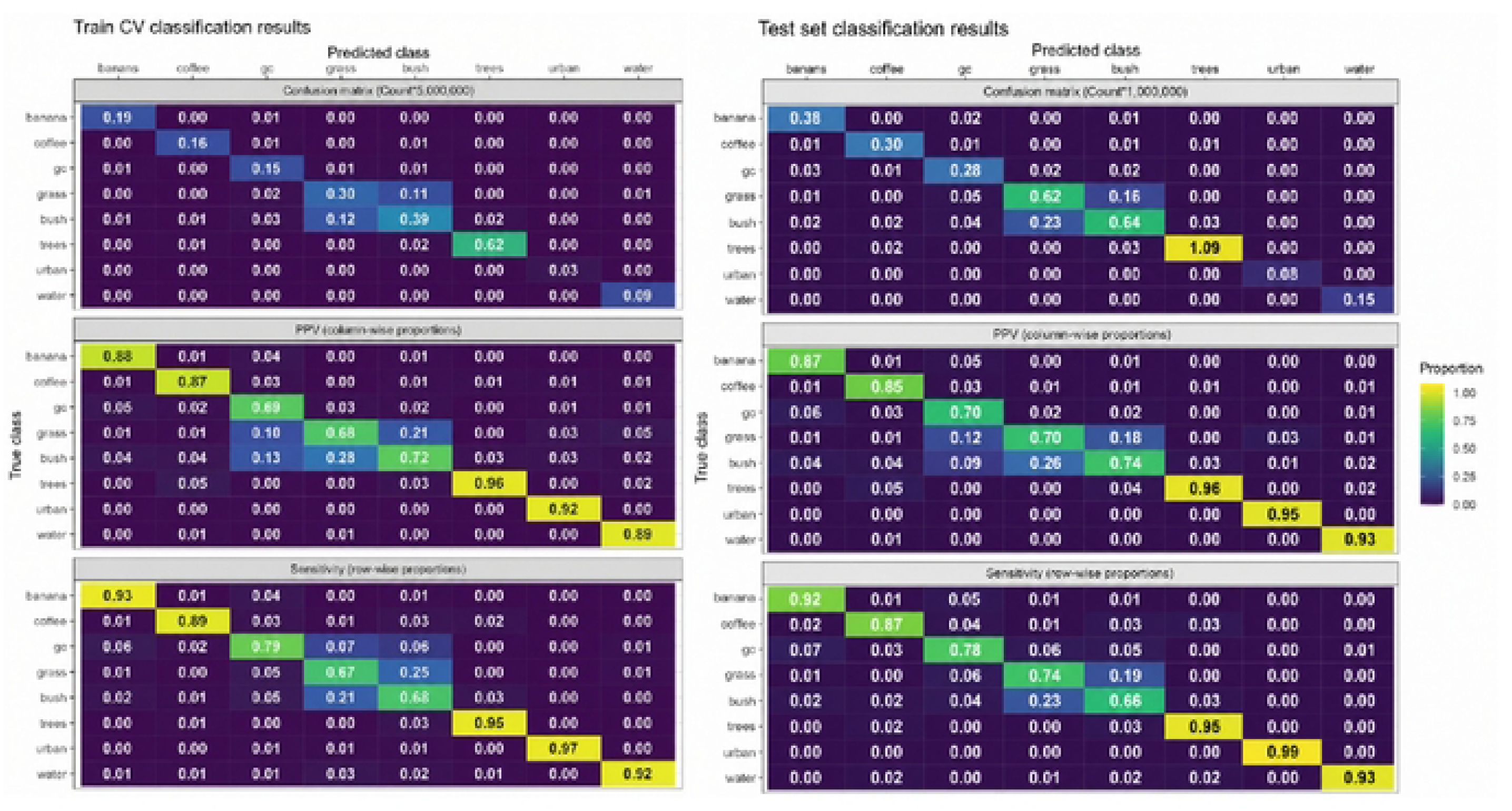
Confusion matrices for cross validation training and test datasets with corresponding row- and column-wise proportions.

**Figure S4.**
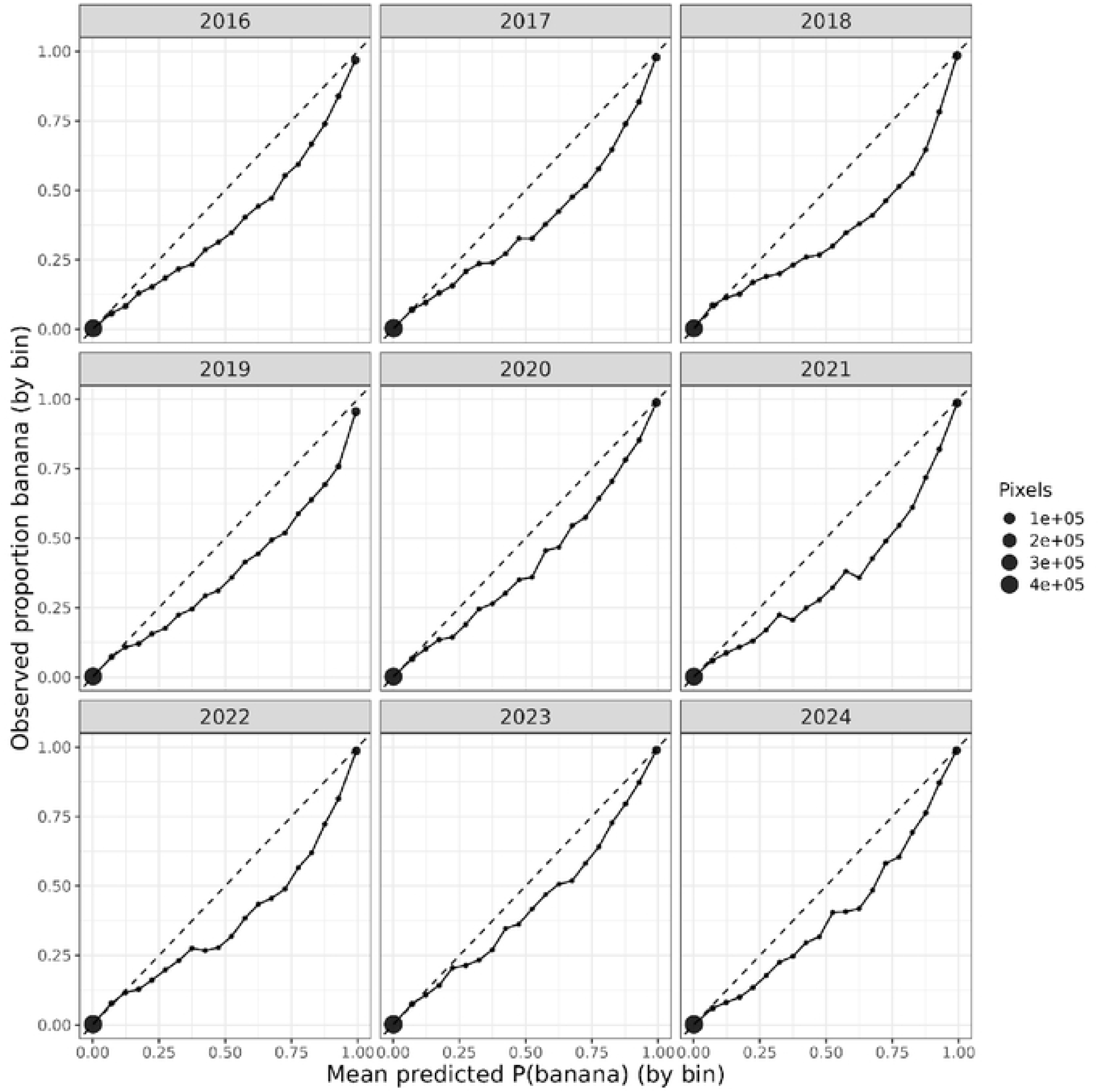
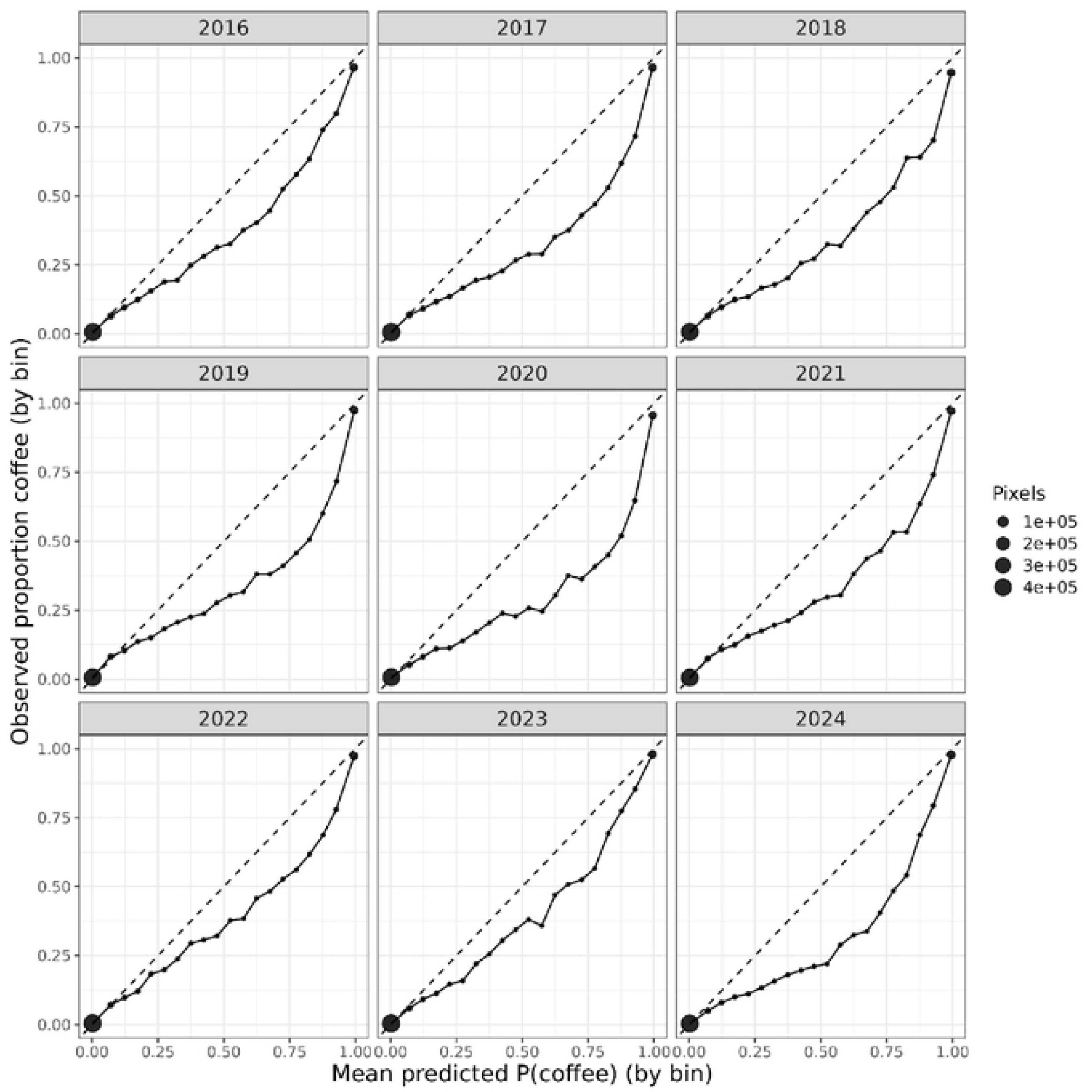
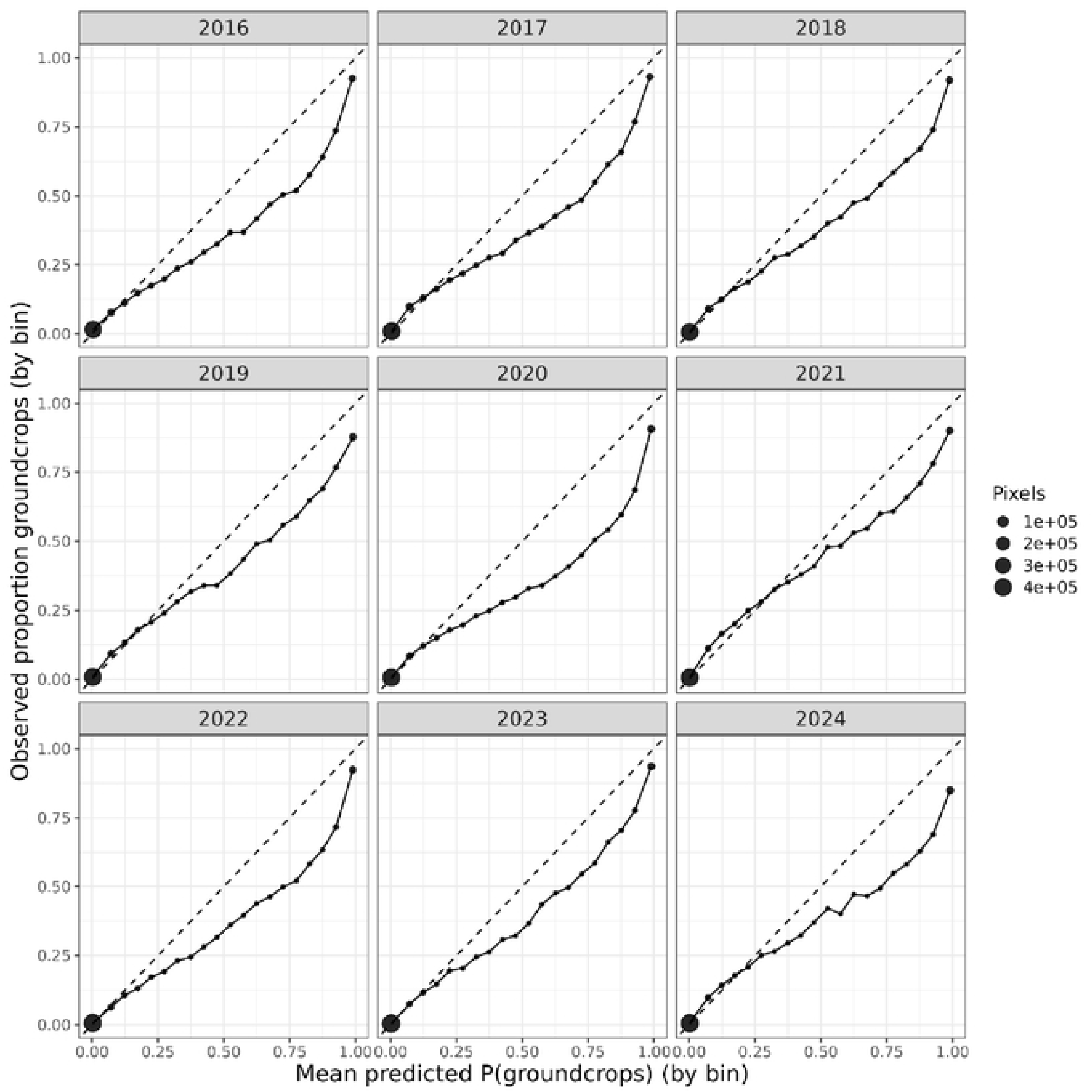
Calibration plots for annual classification of bananas (top), coffee (middle), and ground crops (bottom), visualizing binned predicted probability of crop type (x-axis) and observed proportion belonging to each class (y-axis) by year.

**Figure S5.**
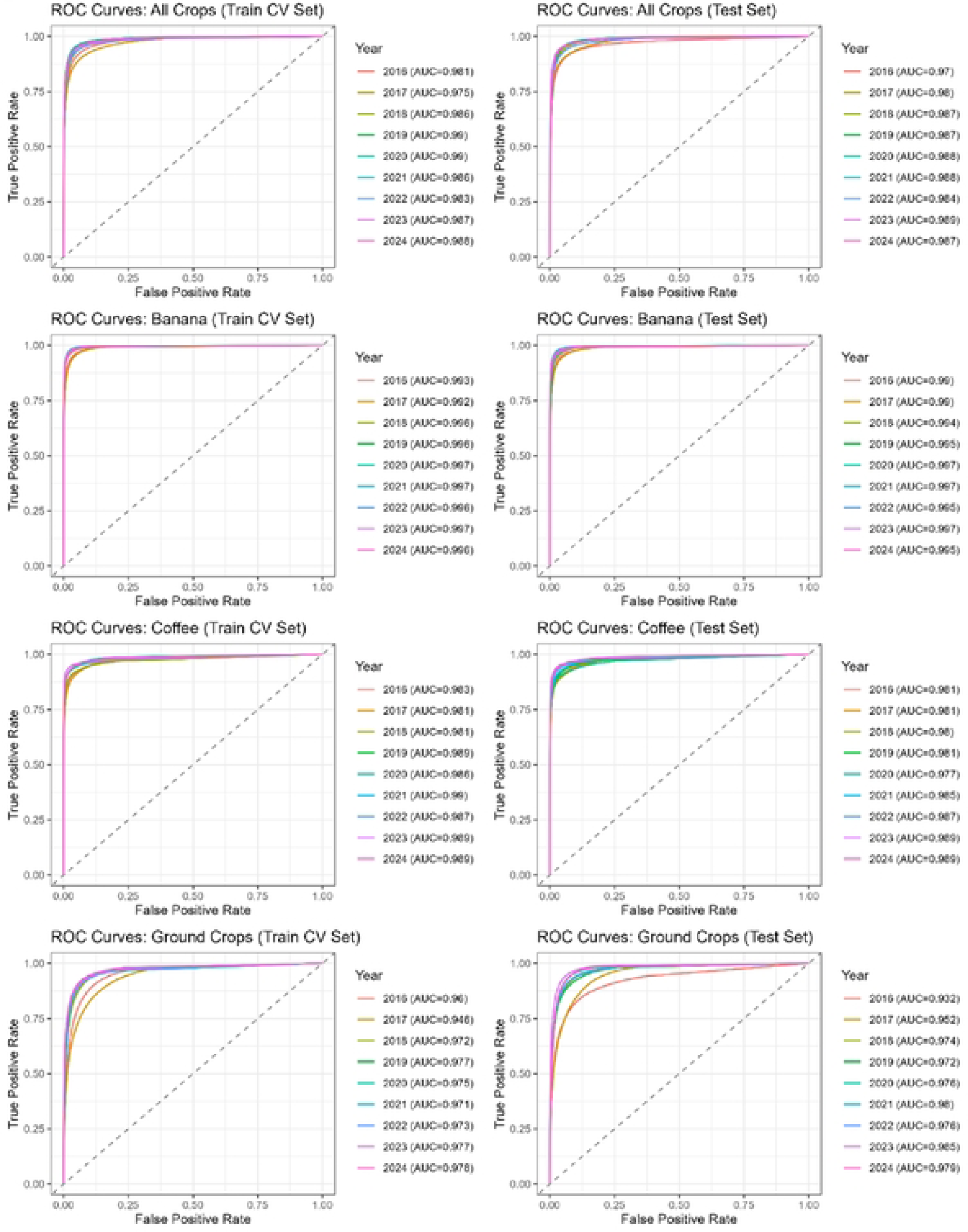
Annual receiver operator curves (ROC) for prediction performance on training and test datasets from 2016-2024 for cumulative crops, and individually for banana, coffee, and ground crops.

**Figure S6.**
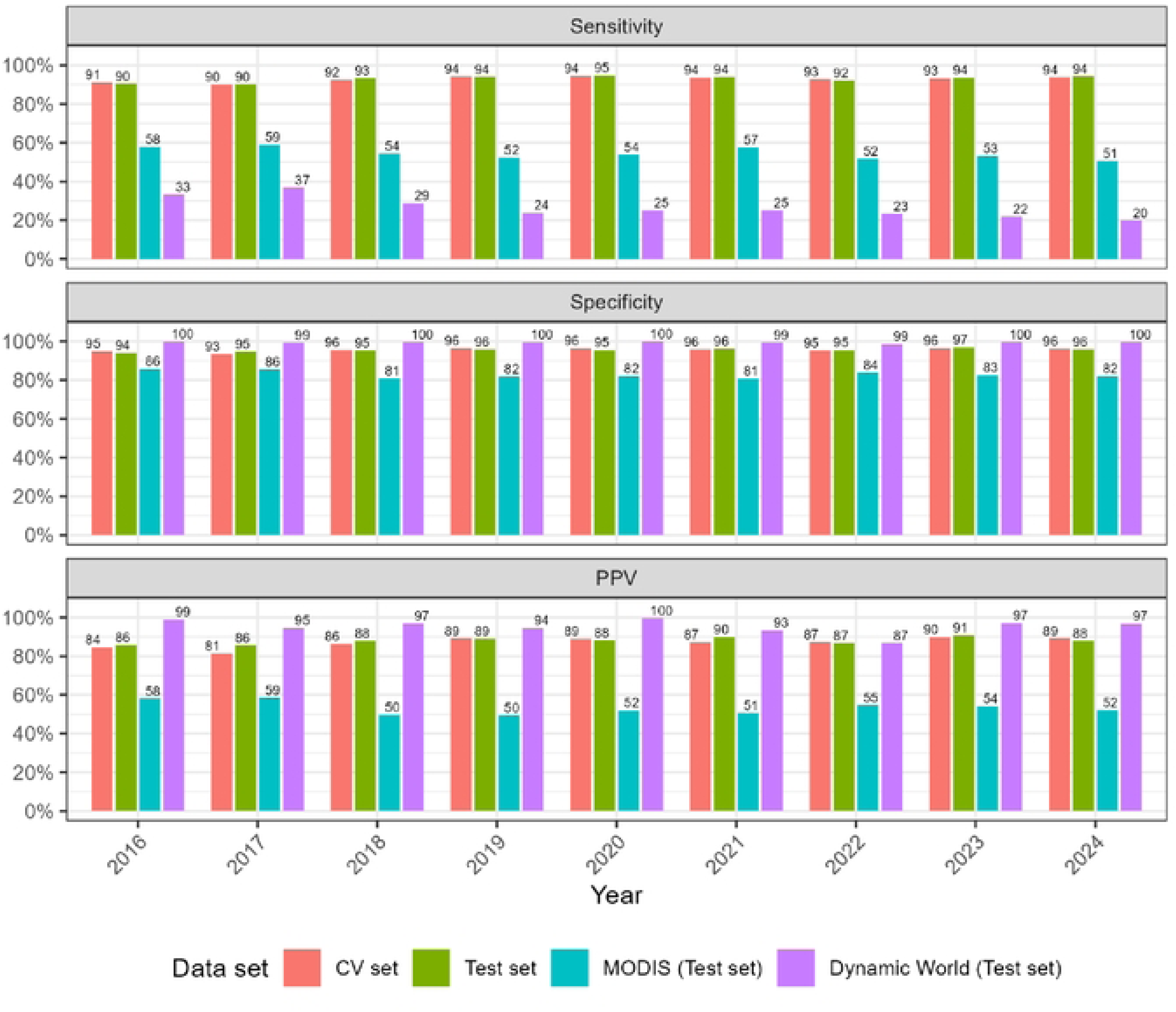
Annual classification performance on pooled crop classes (banana, coffee, and ground crops) resulting from cross validation on the training data set, prediction on the holdout test set, and classification performance of MODIS and Dynamic World datasets on the holdout test set.

**Figure S7.**
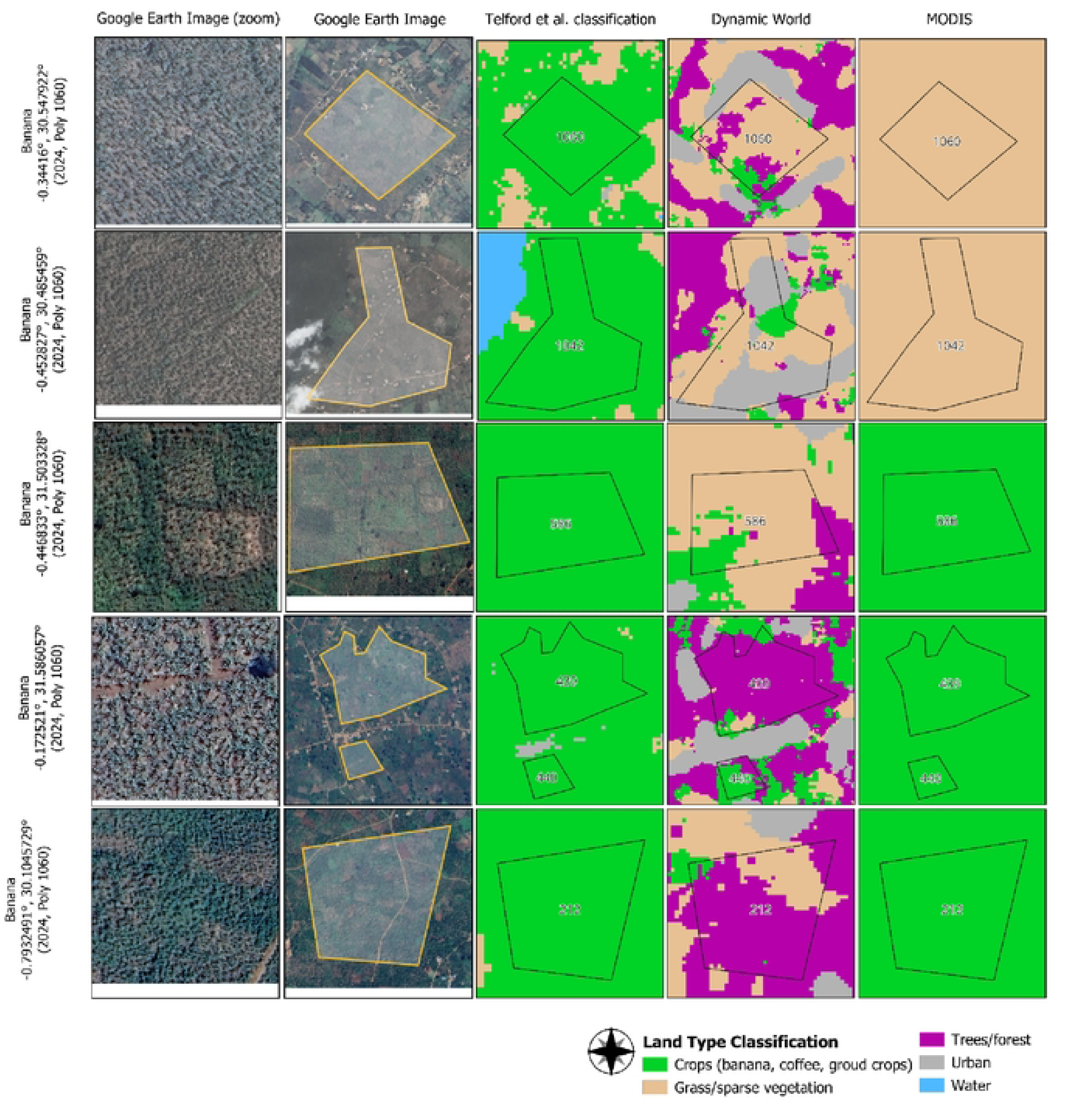
Retrospective evaluation of human-labeled **banana** training polygons with discrepant classifications between Uganda-specific land type classification model in comparison to Dynamic World and MODIS land cover classifications.

**Figure S8.**
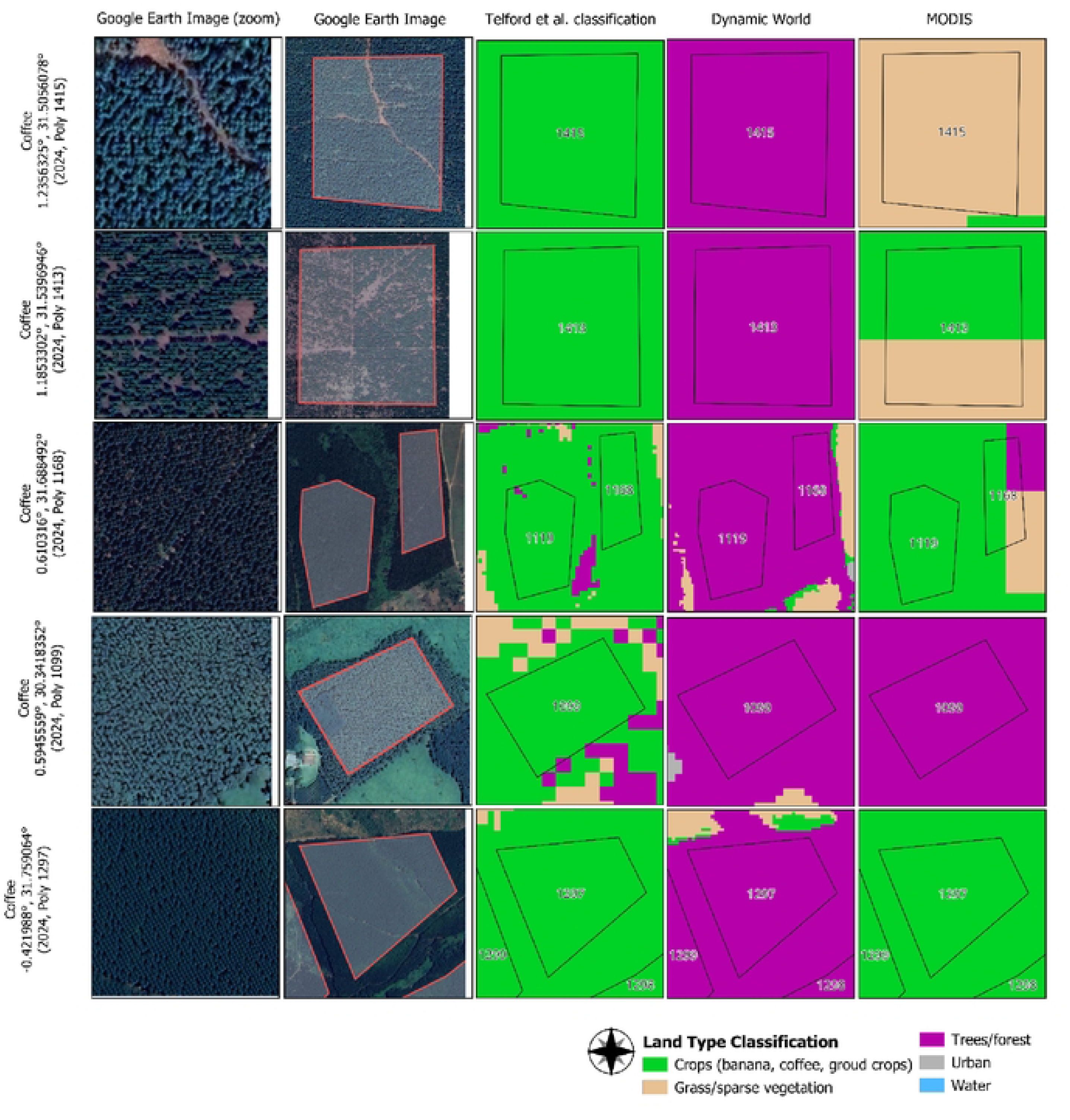
Retrospective evaluation of human-labeled **coffee** training polygons with discrepant classifications between Uganda-specific land type classification model in comparison to Dynamic World and MODIS land cover classifications.

**Figure S9.**
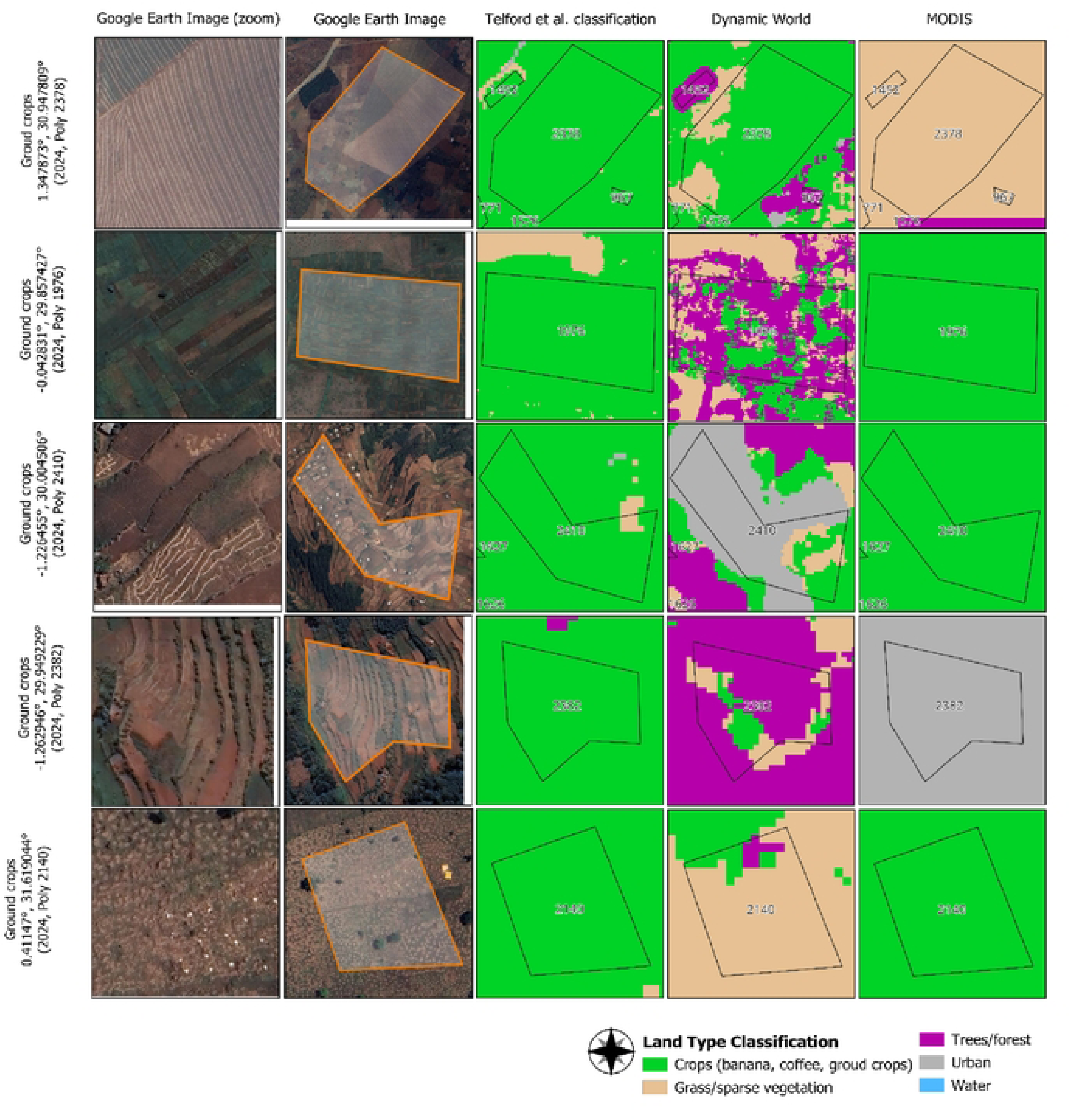
Retrospective evaluation of human-labeled **ground crops** training polygons with discrepant classifications between Uganda-specific land type classification model in comparison to Dynamic World and MODIS land cover classifications.

**Figure S10.**
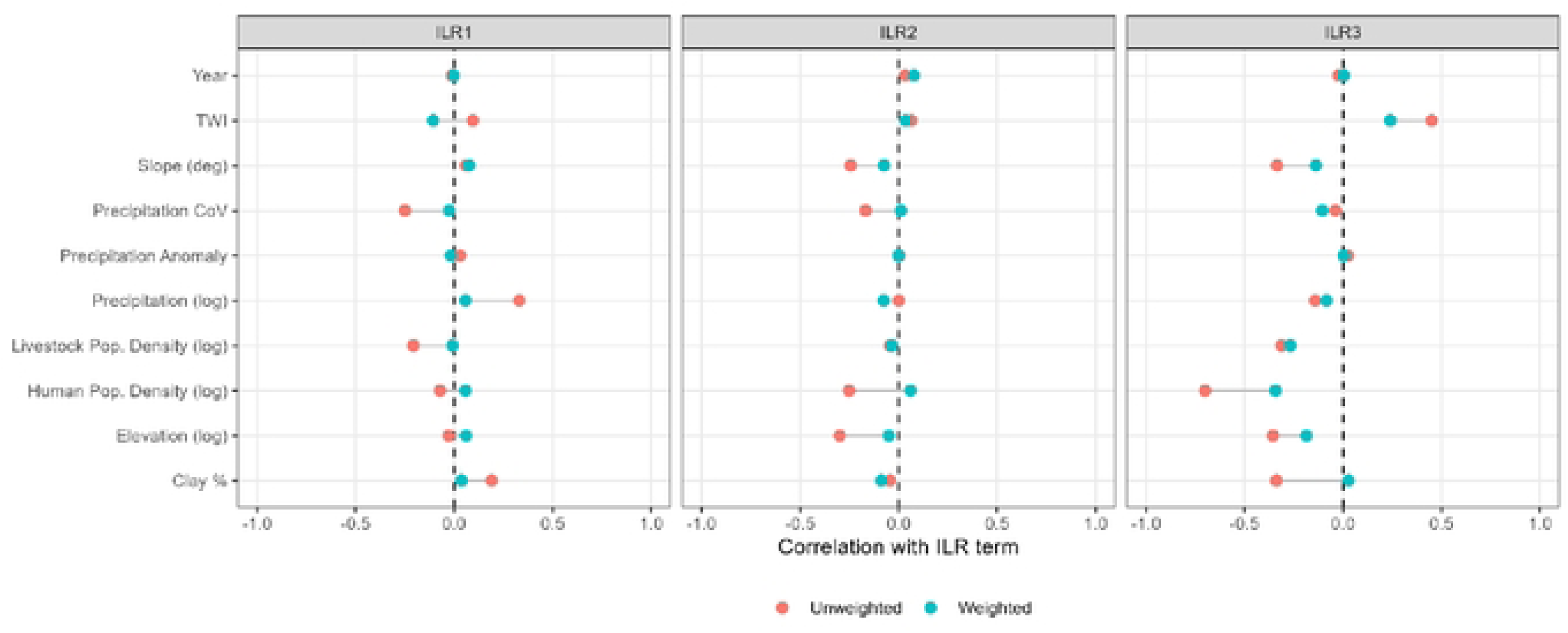
Correlation coefficients for the relationship between confounding variables and the isometric log ratio ([LR) crop land composition terms pre- and post-application of inverse joint density weights estimated from propensity models of ILR terms.

**Figure S11.**
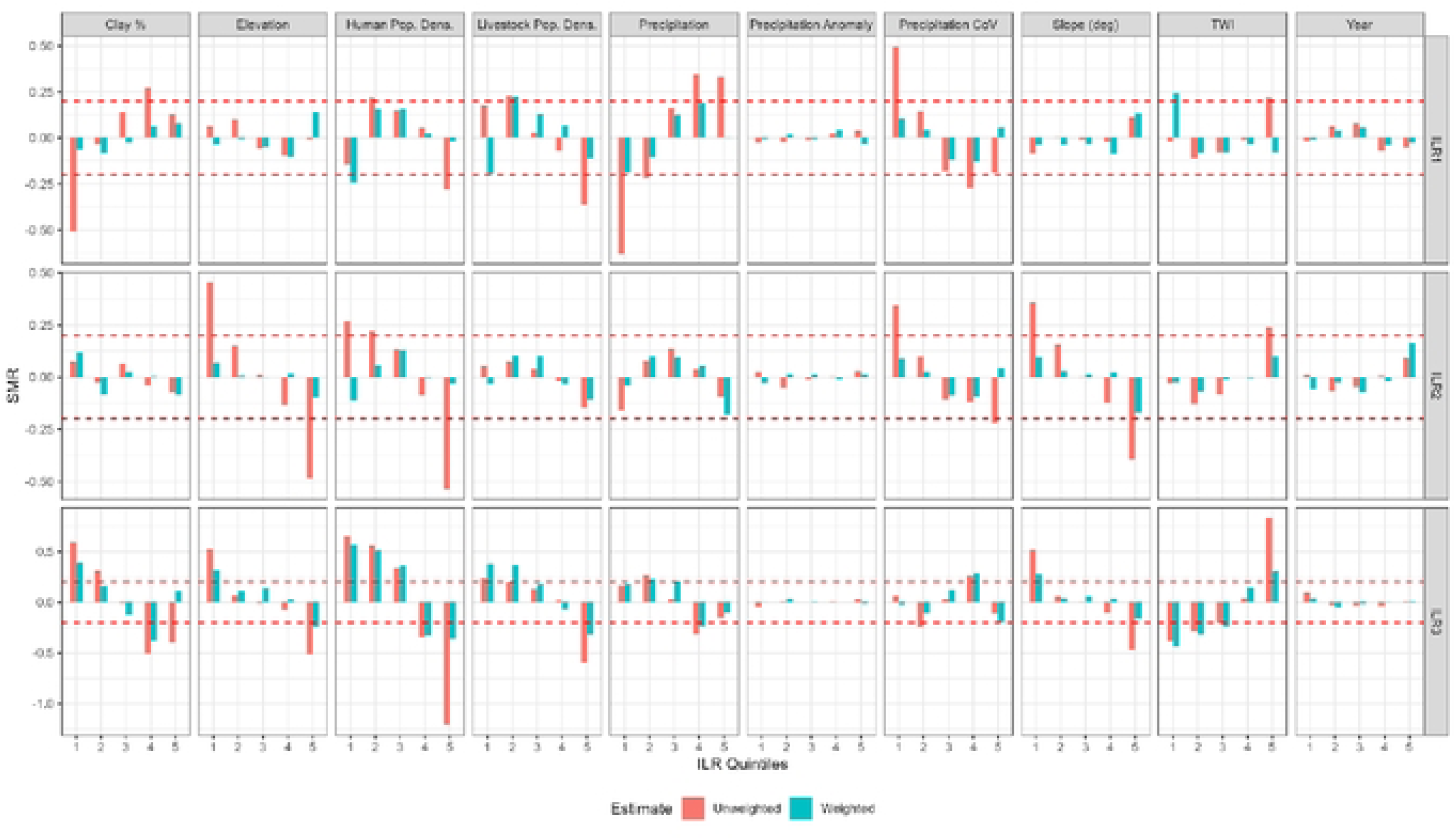
Standardized mean differences (SMDs) in confounding variables across decile levels of crop land composition ILR terms before and after applying stabilized inverse joint density weights.

**Figure S12.**
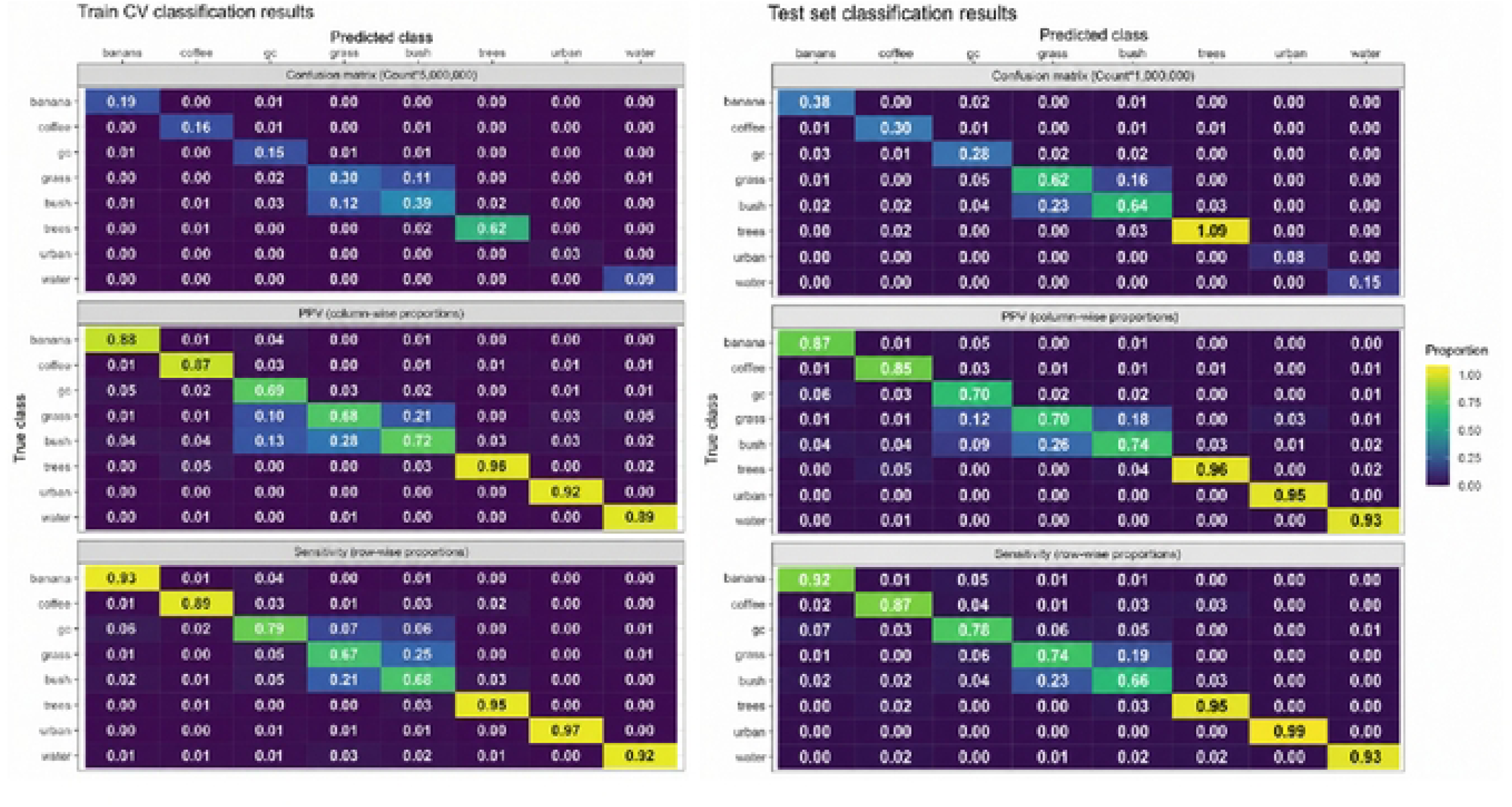
Classification metrics for primary analysis compared sensitivity analysis, which classified land cover based on stochastic sampling within pixel-wise land classification probability distribution.

**Figure S13.**
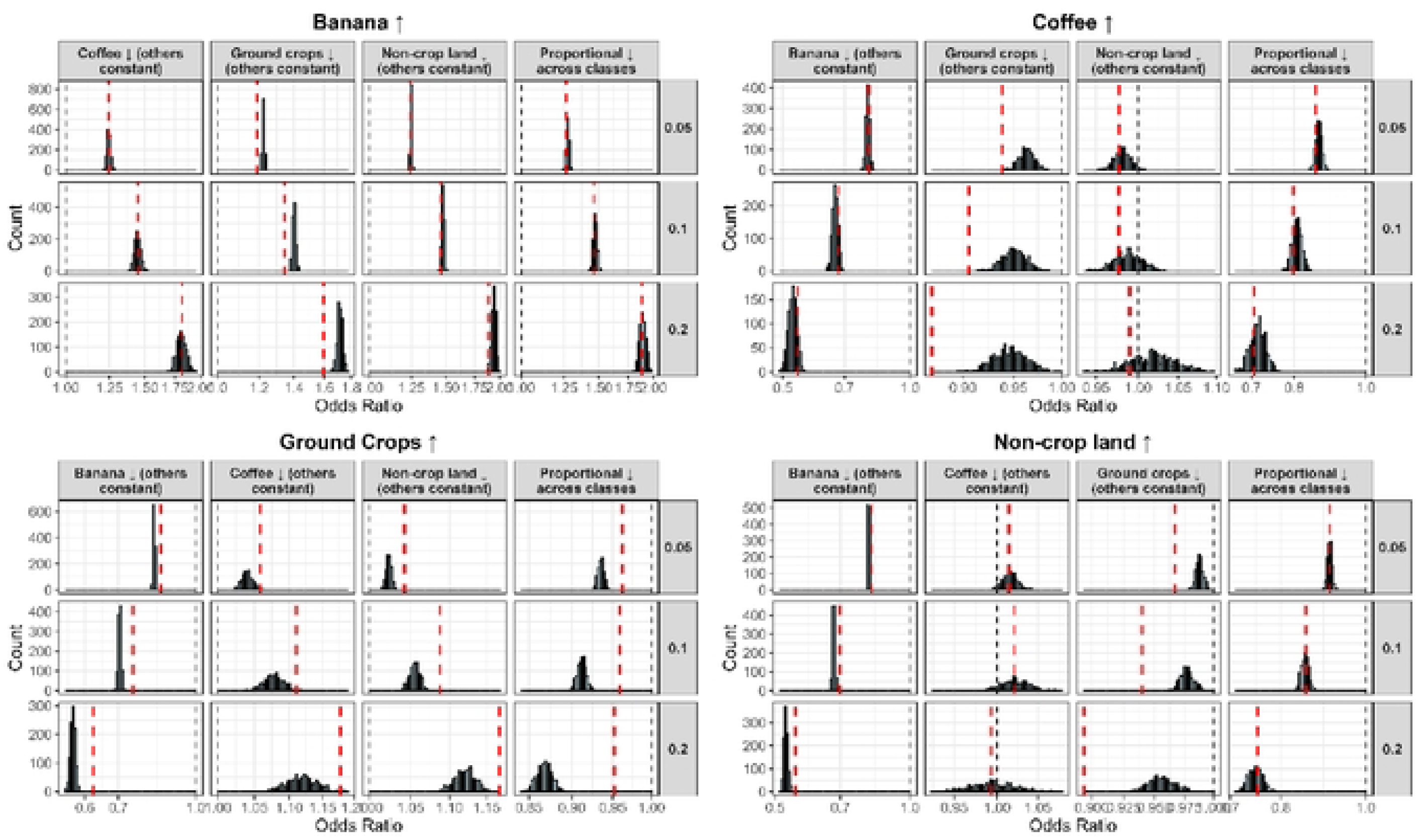
Odds ratio estimates for substitution effects between banana, coffee, ground crops, and non-crop land (columns) across varying substitution magnitudes (rows). The red dotted lines indicate the OR estimate from the main analysis averaged across all low, moderate, and high baseline proportions of the receiving land class. Bars indicate OR bins with a count corresponding to each iteration of the sensitivity analysis (N=JOOO).

**Table S1.** Counts and percent representation of human-labeled training polygons by land-cover class and training dataset period.

| Training Dataset | Class | N Polygons | % Representation |
| --- | --- | --- | --- |
| 2016-2017 |  |  |  |
|  | Banana | 1100 | 8.52% |
|  | Coffee | 769 | 5.95% |
|  | Ground crops | 606 | 4.69% |
|  | Grass | 525 | 4.06% |
|  | Bush | 623 | 4.82% |
|  | Trees | 288 | 2.23% |
|  | Urban | 264 | 2.04% |
|  | Water | 176 | 1.36% |
| 2018-2021 |  |  |  |
|  | Banana | 1094 | 8.47% |
|  | Coffee | 755 | 5.84% |
|  | Ground crops | 463 | 3.58% |
|  | Grass | 610 | 4.72% |
|  | Bush | 565 | 4.37% |
|  | Trees | 287 | 2.22% |
|  | Urban | 250 | 1.94% |
|  | Water | 172 | 1.33% |
| 2022-2024 |  |  |  |
|  | Banana | 1076 | 8.33% |
|  | Coffee | 835 | 6.46% |
|  | Ground crops | 530 | 4.10% |
|  | Grass | 614 | 4.75% |
|  | Bush | 559 | 4.33% |
|  | Trees | 286 | 2.21% |
|  | Urban | 279 | 2.16% |
|  | Water | 192 | 1.49% |

**Table S2.** Collapsed-class confusion matrix for the Uganda-specific land-cover classification model on held-out test pixels across 2016-2024. Non-crop land includes grass, bush, trees, and urban classes; precision is column-wise, recall is row-wise, and cumulative accuracy is the diagonal cumulative proportion.

| True Class | Predicted class |  |  |  |  | Recall** |
| --- | --- | --- | --- | --- | --- | --- |
|  | Banana | Coffee | Ground crops | Non-crops | Water |  |
| Banana | 953463 | 13007 | 56326 | 61624 | 2817 | 87.70% |
| Coffee | 7385 | 817582 | 19860 | 89861 | 7016 | 86.82% |
| Ground crops | 41052 | 30676 | 739989 | 249596 | 4049 | 69.46% |
| Non-crop land | 20213 | 58534 | 115909 | 8101956 | 28925 | 97.31% |
| Water | 588 | 3126 | 4691 | 50416 | 461254 | 88.69% |
| Precision* | 93.23% | 88.59% | 78.99% | 94.72% | 91.51% |  |
| Cumulative Accuracy*** | 92.75% |  |  |  |  |  |

\*Synonymous with positive predicted value or user's accuracy. Calculated as column-wise proportion. \*\*Synonymous with sensitivity or producer's accuracy. Calculated as row-wise proportions. \*\*\*Calculated as diagonal cumulative proportion.

**Table S3.** Mean distributions of primary-analysisand stochastic-sensitivity land-cover compositional proportions among 5 x 5-kilometer grid cells in southwestern Uganda, by RVF outbreak status and overall.

|  | No Outbreak<br>(N=27018) | RVF<br>Outbreak<br>(N=90) | Overall<br>(N=27108) |
| --- | --- | --- | --- |
|  | Mean (SD) | Mean (SD) | Mean (SD) |
| **Land Cover Proportion** |  |  |  |
| Pooled crops | 0.49 (± 0.32) | 0.55 (± 0.30) | 0.49 (± 0.32) |
| Non-crop land | 0.51 (± 0.32) | 0.45 (± 0.30) | 0.51 (± 0.32) |
| Banana | 0.14 (± 0.15) | 0.23 (± 0.19) | 0.14 (± 0.15) |
| Coffee | 0.086 (± 0.097) | 0.099 (± 0.11) | 0.086 (± 0.097) |
| Ground crops | 0.27 (± 0.22) | 0.23 (± 0.21) | 0.27 (± 0.22) |
| **Crop land measures**<br>**(sensitivity analysis)** |  |  |  |
| Pooled crop land | 0.48 (± 0.33) | 0.54 (± 0.30) | 0.48 (± 0.32) |
| Non-crop land | 0.52 (± 0.32) | 0.46 (± 0.30) | 0.52 (± 0.32) |
| Banana crop land | 0.14 (± 0.14) | 0.22 (± 0.18) | 0.14 (± 0.14) |
| Coffee crop land | 0.092 (± 0.095) | 0.10 (± 0.11) | 0.092 (± 0.095) |
| Ground crop land | 0.25 (± 0.19) | 0.22 (± 0.19) | 0.25 (± 0.19) |

## References

1. Prioritizing diseases for research and development in emergency contexts. https://www.who.int/activities/prioritizing-diseases-for-research-and-development-in-emergency-contexts.

2. CDC. About Rift Valley Fever (RVF). Rift Valley Fever https://www.cdc.gov/rift-valley-fever/about/index.html (2024).

3. Linthicum, K. J., Britch, S. C. & Anyamba, A. Rift Valley Fever: An Emerging Mosquito-Borne Disease. Annu. Rev. Entomol. 61, 395–415 (2016).

4. Anyamba, A. et al. Prediction of a Rift Valley fever outbreak. Proc. Natl. Acad. Sci. U.S.A. 106, 955–959 (2009).

5. Turell, M. J. et al. Vector competence of selected African mosquito (Diptera: Culicidae) species for Rift Valley fever virus. Journal of medical entomology 45, 102–108 (2008).

6. Linthicum, K. J., Bailey, C. L., Davies, F. G. & Tucker, C. J. Detection of Rift Valley Fever Viral Activity in Kenya by Satellite Remote Sensing Imagery. Science 235, 1656–1659 (1987).

7. Linthicum, K. J. et al. Towards real-time prediction of Rift Valley fever epidemics in Africa. Preventive Veterinary Medicine 11, 325–334 (1991).

8. Linthicum, K. J. et al. Climate and Satellite Indicators to Forecast Rift Valley Fever Epidemics in Kenya. Science 285, 397–400 (1999).

9. de Glanville, W. A. et al. Inter-epidemic Rift Valley fever virus infection incidence and risks for zoonotic spillover in northern Tanzania. PLoS neglected tropical diseases 16, e0010871 (2022).

10. Kariuki Njenga, M. & Bett, B. Rift Valley Fever Virus—How and Where Virus Is Maintained During Inter-epidemic Periods. Curr Clin Micro Rpt 6, 18–24 (2019).

11. Sumaye, R. D., Geubbels, E., Mbeyela, E. & Berkvens, D. Inter-epidemic transmission of Rift Valley fever in livestock in the Kilombero River Valley, Tanzania: a cross-sectional survey. PLoS Neglected Tropical Diseases 7, e2356 (2013).

12. Sumaye, R. D. et al. Inter-epidemic acquisition of Rift Valley fever virus in humans in Tanzania. PLoS neglected tropical diseases 9, e0003536 (2015).

13. Nyakarahuka, L. et al. Detection of sporadic outbreaks of rift valley fever in Uganda through the National viral hemorrhagic fever surveillance system, 2017–2020. The American Journal of Tropical Medicine and Hygiene 108, 995 (2023).

14. Shoemaker, T. R. et al. First laboratory-confirmed outbreak of human and animal Rift Valley fever virus in Uganda in 48 years. The American Journal of Tropical Medicine and Hygiene 100, 659 (2019).

15. Telford, C., Nyakarahuka, L., Waller, L., Kitron, U. & Shoemaker, T. Geostatistical modeling and prediction of rift valley fever seroprevalence among livestock in Uganda. The American Journal of Tropical Medicine and Hygiene 108, 712 (2023).

16. Nyakarahuka, L. et al. Ten outbreaks of rift valley fever in Uganda 2016-2018: epidemiological and laboratory findings. International Journal of Infectious Diseases 79, 4 (2019).

17. Janko, M. M. et al. The links between agriculture, Anopheles mosquitoes, and malaria risk in children younger than 5 years in the Democratic Republic of the Congo: a population-based, cross-sectional, spatial study. The Lancet Planetary Health 2, e74–e82 (2018).

18. Richards, E. E. et al. The relationship between mosquito abundance and rice field density in the Republic of Korea. Int J Health Geogr 9, 32 (2010).

19. Wang, S., He, X. & Xing, J. Role of agricultural practices in shaping mosquito habitats. Journal of Mosquito Research 14, (2024).

20. Perrin, A., Schaffner, F., Christe, P. & Glaizot, O. Relative effects of urbanisation, deforestation, and agricultural development on mosquito communities. Landsc Ecol 38, 1527–1536 (2023).

21. Muturi, E. J. et al. Mosquito species diversity and abundance in relation to land use in a riceland agroecosystem in Mwea, Kenya. Journal of Vector Ecology 31, 129–137 (2006).

22. UGANDA BUREAU OF STATISTICS. Annual Agricultural Survey (AAS) 2022 – Statistical Release. https://www.ubos.org/wp-content/uploads/publications/05_2022Uganda_UBOS_StatRelease_AAS2019-Final.pdf.

23. Haddow, A. J. The mosquitoes of Bwamba County, Uganda. II.—Biting activity with special reference to the influence of microclimate. Bulletin of Entomological Research 36, 33–73 (1946).

24. Haddow, A. J. The mosquitoes of Bwamba County, Uganda. VI.—Mosquito breeding in plant axils. Bulletin of Entomological Research 39, 185–212 (1948).

25. Center for International Earth Science Information Network - CIESIN - Columbia University. 2018. Gridded Population of the World, Version 4 (GPWv4): Population Density, Revision 11. Palisades, NY: NASA Socioeconomic Data and Applications Center (SEDAC). 10.7927/H49C6VHW.

26. Shoemaker, T. R. et al. Impact of enhanced viral haemorrhagic fever surveillance on outbreak detection and response in Uganda. The Lancet Infectious Diseases 18, 373–375 (2018).

27. Bird, B. H., Bawiec, D. A., Ksiazek, T. G., Shoemaker, T. R. & Nichol, S. T. Highly Sensitive and Broadly Reactive Quantitative Reverse Transcription-PCR Assay for High-Throughput Detection of Rift Valley Fever Virus. J Clin Microbiol 45, 3506–3513 (2007).

28. Gorelick, N. et al. Google Earth Engine: Planetary-scale geospatial analysis for everyone. Remote sensing of Environment 202, 18–27 (2017).

29. European Space Agency. Sentinel-2 Level-1C data. Copernicus Programme (2015).

30. Goward, S. N., Markham, B., Dye, D. G., Dulaney, W. & Yang, J. Normalized difference vegetation index measurements from the advanced very high resolution radiometer. Remote sensing of environment 35, 257–277 (1991).

31. Merton, R. Monitoring community hysteresis using spectral shift analysis and the red-edge vegetation stress index. in Proceedings of the Seventh Annual JPL Airborne Earth Science Workshop 12–16 (JPL Pasadena, CA, USA, 1998).

32. R Core Team. R: A language and environment for statistical computing. R Foundation for Statistical Computing.

33. Kuhn, M. Building predictive models in R using the caret package. Journal of statistical software 28, 1–26 (2008).

34. Westreich, D. et al. Causal impact: epidemiological approaches for a public health of consequence. American journal of public health 106, 1011 (2016).

35. Egozcue, J. J., Pawlowsky-Glahn, V., Mateu-Figueras, G. & Barceló-Vidal, C. Isometric Logratio Transformations for Compositional Data Analysis. Mathematical Geology 35, 279–300 (2003).

36. Aitchison, J. The Statistical Analysis of Compositional Data. Journal of the Royal Statistical Society Series B: Statistical Methodology 44, 139–160 (1982).

37. Arnold, K. F., Berrie, L., Tennant, P. W. & Gilthorpe, M. S. A causal inference perspective on the analysis of compositional data. International journal of epidemiology 49, 1307–1313 (2020).

38. Dumuid, D. et al. Compositional data analysis for physical activity, sedentary time and sleep research. Stat Methods Med Res 27, 3726–3738 (2018).

39. Snowden, J. M., Rose, S. & Mortimer, K. M. Implementation of G-computation on a simulated data set: demonstration of a causal inference technique. American journal of epidemiology 173, 731–738 (2011).

40. Williams, J. R. & Crespi, C. M. Causal inference for multiple continuous exposures via the multivariate generalized propensity score. *arXiv preprint arXiv:2008.13767* https://arxiv.org/abs/2008.13767 (2020).

41. Van der Laan, M. J., Polley, E. C. & Hubbard, A. E. Super learner. https://biostats.bepress.com/ucbbiostat/paper222/ (2007).

42. Kainz, K. et al. Improving Causal Inference: Recommendations for Covariate Selection and Balance in Propensity Score Methods. Journal of the Society for Social Work and Research 8, 279–303 (2017).

43. Austin, P. C. Balance diagnostics for comparing the distribution of baseline covariates between treatment groups in propensity-score matched samples. Statistics in Medicine 28, 3083–3107 (2009).

44. Lutwama, J. J. Seasonal Distribution of Aedes (Stegomyia) simpsoni (Theobald) Immatures in Plant Axils of Araceae and Musaceae in Uganda. Int J Trop Insect Sci 18, 105–112 (1998).

45. Karungi, J. et al. Elevation and cropping system as drivers of microclimate and abundance of soil macrofauna in coffee farmlands in mountainous ecologies. Applied Soil Ecology 132, 126–134 (2018).

46. Geissbühler, Y. et al. Microbial larvicide application by a large-scale, community-based program reduces malaria infection prevalence in urban Dar es Salaam, Tanzania. PloS one 4, e5107 (2009).

47. Mazigo, H. D., Mboera, L. E. G., Rumisha, S. F. & Kweka, E. J. Malaria mosquito control in rice paddy farms using biolarvicide mixed with fertilizer in Tanzania: semi-field experiments. Malar J 18, 226 (2019).

48. Nyakarahuka, L. et al. Prevalence and risk factors of Rift Valley fever in humans and animals from Kabale district in Southwestern Uganda, 2016. PLoS neglected tropical diseases 12, e0006412 (2018).

49. Tumusiime, D. et al. Participatory survey of risk factors and pathways for Rift Valley fever in pastoral and agropastoral communities of Uganda. Preventive veterinary medicine 221, 106071 (2023).

